# Spatiotemporal transcriptomic maps of mouse intracerebral hemorrhage at single-cell resolution

**DOI:** 10.1101/2024.05.23.595245

**Authors:** Rong Xiang, Junmin Wang, Mingyue Wang, Zhan Chen, Jin Tao, Ruoqi Ding, ZhenCheng Tu, Shaoshuai Wang, Tao Yang, Jing Chen, Zihan Jia, Qinfeng Peng, Xueping Li, Xinru Zhang, Shuai Chen, Nannan Cheng, Mengke Zhao, Jiaxin Li, Qidi Xue, Chao Jiang, Yolanda de Pablo, Ulrika Wilhelmsson, Marcela Pekna, Milos Pekny, Nicholas Mitsios, Chuanyu Liu, Xun Xu, Xiaochong Fan, Jan Mulder, Xuemei Chen, Longqi Liu, Jian Wang

## Abstract

Intracerebral hemorrhage (ICH) is a severe and widespread disease that results in high mortality and morbidity. Despite significant advances made in clinical and preclinical research, the prognosis of ICH remains poor due to an incomplete understanding of the complex pathological responses. To address this challenge, we generated single-cell resolution spatiotemporal transcriptomic maps of mouse brain 1, 3, 7, 14, and 28 days after ICH. Our analysis revealed the cellular constituents and their dynamic responses following ICH. We found changes in gene expression that are indicative of active phagocytosis and lipid processing in the lesion core and identified genes associated with brain repair at the lesion border. Persistent lipid accumulation culminated in the phenotypic transition of macrophages into foam cells. Furthermore, our results demonstrate that ICH could stimulate neural stem cells in the subventricular zone, thereby enhancing neurogenesis in this niche. This study provides a spatiotemporal molecular atlas of mouse ICH that advances the understanding of both local and global responses of brain cells to ICH and offers a valuable resource that can aid the development of novel therapies for this devastating condition.

## Introduction

Intracerebral hemorrhage (ICH) is a type of stroke that accounts for 15-20% of all strokes. It is a challenging condition with a high mortality rate of up to 50%^1,2^. Although minimally invasive surgical techniques for hematoma evacuation may be helpful in some instances^3,4^, the prognosis for ICH patients remains unfavorable due to the complex interplay of primary and secondary injury mechanisms that occur in response to ICH^5^.

Brain tissue damage after ICH is due to the initial bleeding, hematoma expansion, brain edema, and damage-associated events such as hemoglobin released by red blood cells. Microglia respond within minutes after injury, disruption of blood-brain barrier and the onset of brain edema^6–9^. The persistence of neuroinflammation and immune responses determines the severity of secondary brain injury, which contributes to a poor prognosis for ICH patients^10,11^. Pro-inflammatory factors, released by microglia attract and activate immune cells from the periphery leading to a series of inflammatory reactions. These early inflammatory responses are essential for hematoma clearance^8,12^, persistent inflammation can, however, compromise neuronal survival^13,14^. Balancing the beneficial and detrimental effects of inflammation in treating ICH remains challenging. ICH additionally triggers endogenous anti-inflammatory, anti-excitatory, antioxidant, and neurotrophic responses that are potentially neuroprotective^15^. Thus, ICH activates a cascade of events that involve a complex interplay of cellular responses, molecular signaling pathways, and dynamic microenvironmental changes^1^.

Most studies on ICH have focused on specific molecules^12,16,17^ and have not been able to address the complexity of cellular and molecular interactions triggered by ICH. In particular, the cellular interactions in the perihematomal region that determine ICH prognosis are poorly understood^18,19^. Spatial transcriptomics is a powerful and unbiased tool that enables the identification of cellular and molecular alterations in the hematoma vicinity as well as in more remote brain regions. To investigate cellular responses, molecular signaling, and dynamic changes at different time points after ICH in the striatum, our study employed a sub-cell resolution spatial transcriptomic technology, Stereo-seq^20^. We observed spatially graded gene expression in and around the hematoma, indicating a distinct molecular response to ICH across different spatial domains and cell-types. Furthermore, ICH significantly altered gene expression in more distal brain regions, where it facilitated the transition of dormant neural stem cells into an active state and, beyond day 7 induced the activation of immediate early genes (IEGs) in the cortex. Our spatial transcriptomic data are available for interactive exploration at (https://db.cngb.org/stomics/stmich/), and we believe they serve as a foundational resource for future research in this area. Our findings provide novel insights into the spatial and temporal pathophysiological processes triggered by ICH and underscore the importance of continued research in this field.

## Results

### Spatiotemporal transcriptomic analysis of ICH

To investigate the changes in gene expression following ICH in terms of temporal and spatial dynamics, we induced ICH in mice by collagenase injection into the striatum^21,22^. This ICH model mimics acute cerebrovascular injury and produces bleeding that is reproducible in terms of location and size^23,24^. As in **Figure 1A**, bleeding was mainly located within the striatum, confirming the modeling accuracy. We have assessed the neurological deficits of ICH mice compared to controls on days 1, 3, 7, 14, and 21 following injury. The results were consistent with our previous reports^12,22^, showing that they had significant neurological deficits at the first four time points (**Figure 1B**). In addition, we observed a notable increase in brain water content in the ipsilateral hemisphere on day 3 after ICH compared to the naive group (80.63±2.85% vs. 77.60±0.49%, *P*<0.05, n=5-6) **(Figure 1C)**. However, we found no difference in brain water content between the two groups on the contralateral side (77.91±0.83% vs.77.88± 0.61%) or cerebellum (75.96 ± 1.21% vs. 75.91±1.93%, n=5-6). Crystal Violet (CV)/Luxol Fast Blue (LFB)-staining showed a typical lesion (volume: 8.77±1.68 mm^3^) and a decrease in the normal myelin area around the lesion compared to the corresponding brain area in the naive group (26.58±6.85% vs. 53.78±11.63%, n=6) on day 3 after ICH **(Figure 1D-1F).** We then sampled all brain sections with selected intervals around the injection site and monitored the extent of the lesion and selected sections with signs of bleeding. Using spatial transcriptomic analysis, we mapped the gene expression changes at day 1, 3, 7, 14, and 28 post-ICH. We have employed Stereo-seq to capture transcripts on these sections at the subcellular level. This technique has enabled us to analyze global gene expression and cellular changes in the mouse brain at different timepoints following ICH **(Figure 1G)**.

**Figure 1.**
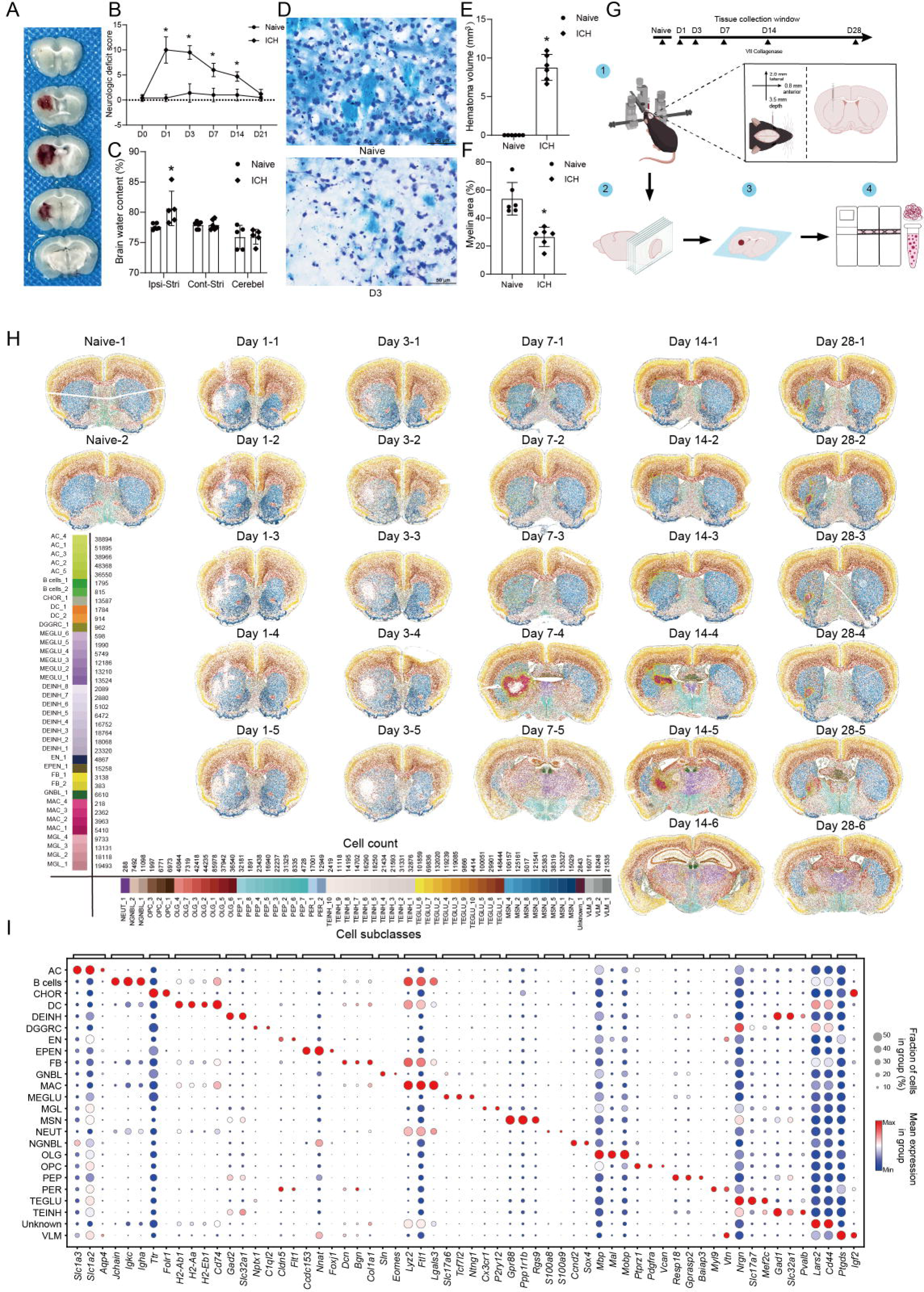
Spatiotemporal transcriptomic map of mouse intracerebral hemorrhage (ICH) at single-cell resolution. (A) Representative fresh brain sections with red areas indicating hematomas in mice on day 3 follwoing ICH. (B) Assessment of neurological deficits. The mice had apparent neurologic deficits on days 1, 3, 7 and 14 after ICH. n=10 mice/group. \**P* <0.05 vs. the naïve group (repeated measures ANOVA followed by Bonferroni’s post hoc test). (C) Determination of brain water content. The brain water content was significantly increased in the ipsilateral hemisphere of mice on day 3 after ICH, compared to the naïve animals. n=5-6 mice/group. \**P*<0.05 vs. the naïve group (t-test). Cont-Stri: contralateral striatum; Ipsi-Stri: ipsilateral striatum; Cerebel: cerebellum. (D) Representative CV/ LFB-stained brain sections from the ICH and naïve groups (Scale bar = 50 μm). CV: Crystal Violet, LFB: Luxol fast blue. (E) Measurement of hematoma volume.The hematoma volume was significantly larger in the ICH than in the naïve group. n=6 mice/group. \**P*<0.05 vs. the naïve group (t-test). (F) Measurement of normal myelin area.There was a significant decrease in the myelinated areas in ICH compared to naïve groups on day 3. n=6 mice/group. \**P*<0.05 vs. the naïve group (t-test). (G) Outline of the main steps of the experimental procedure. 1. Construction of the striatal ICH model; 2. Collection of brain slices; 3. Attachment of tissue sections to the surface of the Stereo-seq capture chip; and 4. Preparation of DNA libraries and subsequent sequencing. (H) Spatiotemporal visualization of cells from all time-points after ICH in mice. Each dot represents a cell. Cells are colored according to cell subclasses assignment. (I) Dot plot of the main signature genes for all cell types. Average marker gene expression is displayed after z-score transformation. Abbreviation of cell types: DGGRC, dentate gyrus granule neurons; MEGLU, dimesencephalon excitatory neurons; DEINH, dimesencephalon inhibitory neurons; GNBL, glutamatergic neuroblasts; NGNBL, non-glutamatergic neuroblasts; PEP, peptidergic neurons; TEINH, telencephalon inhibitory interneurons; TEGLU, telencephalon projecting excitatory neurons; MSN, telencephalon projecting inhibitory neurons; MAC, macrophages; NEUT, neutrophils; DC, dendritic cells; AC, astrocytes; MGL, microglia; OPC, oligodendrocyte precursor cells; OLG, oligodendrocytes; EPEN, ependymal cells; CHOR, choroid plexus epithelial cells; EN, endothelial cells; PER, pericytes; FB, fibroblasts; VLM, vascular and leptomeningeal cells.

To identify the location of individual cells in the tissue section, we used DNA stains to visualize the location of cell nuclei. Then, we employed the cellpose algorithm to segment individual nuclei^25^, and group the transcripts from each nucleus and its surrounding region (≈6 µm). This approach allowed us to attribute transcripts to individual cellular compartments and achieve spatial transcriptomics at a single-cell resolution **(Figure S1A, STAR methods)**. Differential spatial distribution was observed between nucleus-localized transcripts (*Malat1* and *Neat1*) and perinuclear compartment mitochondrial transcripts within individual cells**(Figure S1B)**. After quality control **(Figure S1C)**, we obtained spatial transcriptomics datasets for 2,628,392 cells, averaging 611 genes per cell (**Table S1**).

We have employed a two-tiered annotation process to create an atlas of cell-type responses in the mouse brain following ICH **(Figure S1C)**. In the first step, we classified the main cell types while, in the second step, we classified the subclasses or cell states. We conducted batch correction of Stereo-seq data with scVI^26^ and performed initial cell classification using the semi-supervised algorithm scANVI^27^. This approach labels query data by querying reference data^28^. After automated classification, we manually identified 23 main cell types in the Stereo-seq data using curated marker genes from the literature **(Table S2)**. These cell types included 9 neuronal, 4 immune, 5 glial, and 5 vascular cell clusters **(Figure 1H)**. The neuronal clusters included dentate gyrus granule neurons (DGGRC), diencephalon and mesencephalon excitatory neurons (MEGLU), diencephalon and mesencephalon inhibitory neurons (DEINH), glutamatergic neuroblasts (GNBL), non-glutamatergic neuroblasts (NGNBL), peptidergic neurons (PEP), telencephalon inhibitory interneurons (TEINH), telencephalon projecting excitatory neurons (TEGLU), and telencephalon projecting inhibitory neurons (MSN). The immune clusters included macrophages (MAC), B cells, neutrophils (NEUT), and dendritic cells (DC). The glial clusters included astrocytes (AC), microglia (MGL), oligodendrocyte precursor cells (OPC), oligodendrocytes (OLG), and ependymal cells (EPEN). The vascular clusters included choroid plexus epithelial cells (CHOR), endothelial cells (EN), pericytes (PER), fibroblasts (FB), and vascular and leptomeningeal cells (VLM). We manually annotated these main clusters with corresponding canonical marker genes, indicating a robust annotation result (**Figure 1I and S1D**). We also identified a not yet annotated cell type that we named "Unknown," which showed high expression of *Lars2* and *Cd44*. Further clustering and manual annotation within the main clusters resulted in identifying 92 cell subclasses, comprising 54 neuronal, 20 glial, 11 immune, and 7 vascular cell subclasses **(Figure 1H)**. The subclassification enabled us to determine the spatial heterogeneity and gene expression patterns for each cell type **(Figure S2, Table S3)**. For instance, the different subclasses of TEGLU were distributed across various cortical regions, whereas distinct subclasses and markers of astrocytes and microglia were found in different brain regions **(Figure S2A, S2E and S2L).**

Based on the available data, we initially evaluated the activities of the classical signaling pathways using PROGENy^29,30^(**Figure S3A, STAR methods**). Within 24h after ICH, we observed sporadic spots of JAK-STAT signaling activity in both the lesion and the vascular pia meningeal area. This signaling activity likely contributed to the inflammatory response and astrocyte activation following ICH^31^. Furthermore, the lesion exhibited predominant pro-inflammatory TNFα signaling, alongside the presence of counter acting anti-inflammatory TGFβ signaling. In addition, we noticed increased activity in WNT signaling and EGFR signaling pathways after day 7, which indicates the initiation of tissue remodeling and repair. In summary, the spatial data depicted the typical pathologic processes and changes in cell types within the mouse brain after ICH.

### Spatiotemporal distribution of ICH-associated cell subclasses

ICH triggers a complex cascade of cellular responses within the brain, forming a unique lesion microenvironment. Despite recognizing the participation of specific cell types in lesion formation^32^, the spatiotemporal dynamics of the lesion microenvironment are partially understood.

We used ClusterMap^33^ to build a molecularly defined tissue region map to investigate whether unique lesion tissue domains are formed after ICH in our data (**See Methods**). Overall, these tissue regions exhibited good alignment with anatomically defined regions (**Figure S4A**). For instance, the molecularly defined cortical regions displayed the same layering as the anatomical layers of the cerebral cortex (L2/3, L4, L5, L6). However, we discovered five distinct areas around the bleeding site that are absent in control brains (**Figure S4B**). Three of these regions had cell compositions similar to those in the unaffected striatum, and as such, we named them "Injured_STR_1", "Injured_STR_2", and "Injured_STR_3" (**Figure 2A**). The other two areas, which we have named "Lesion_1" and "Lesion_2", formed the bleeding core that could be identified at five time-points following ICH and had cell compositions distinct from those of the naive striatum. We also examined the cell types in the lesion over time **(Figure 2B)**. We selected brain sections from various time points, including naive (section 2), day 1 (section 3), day 3 (section 3), day 7 (section 4), day 14 (section 4), and day 28 (section 3), based on lesion sizes. Our results showed that the lesion region primarily comprised astrocytes, macrophages, oligodendrocytes, fibroblasts, MSNs, and peripheral immune cells such as neutrophils, B cells, and dendritic cells.

**Figure 2.**
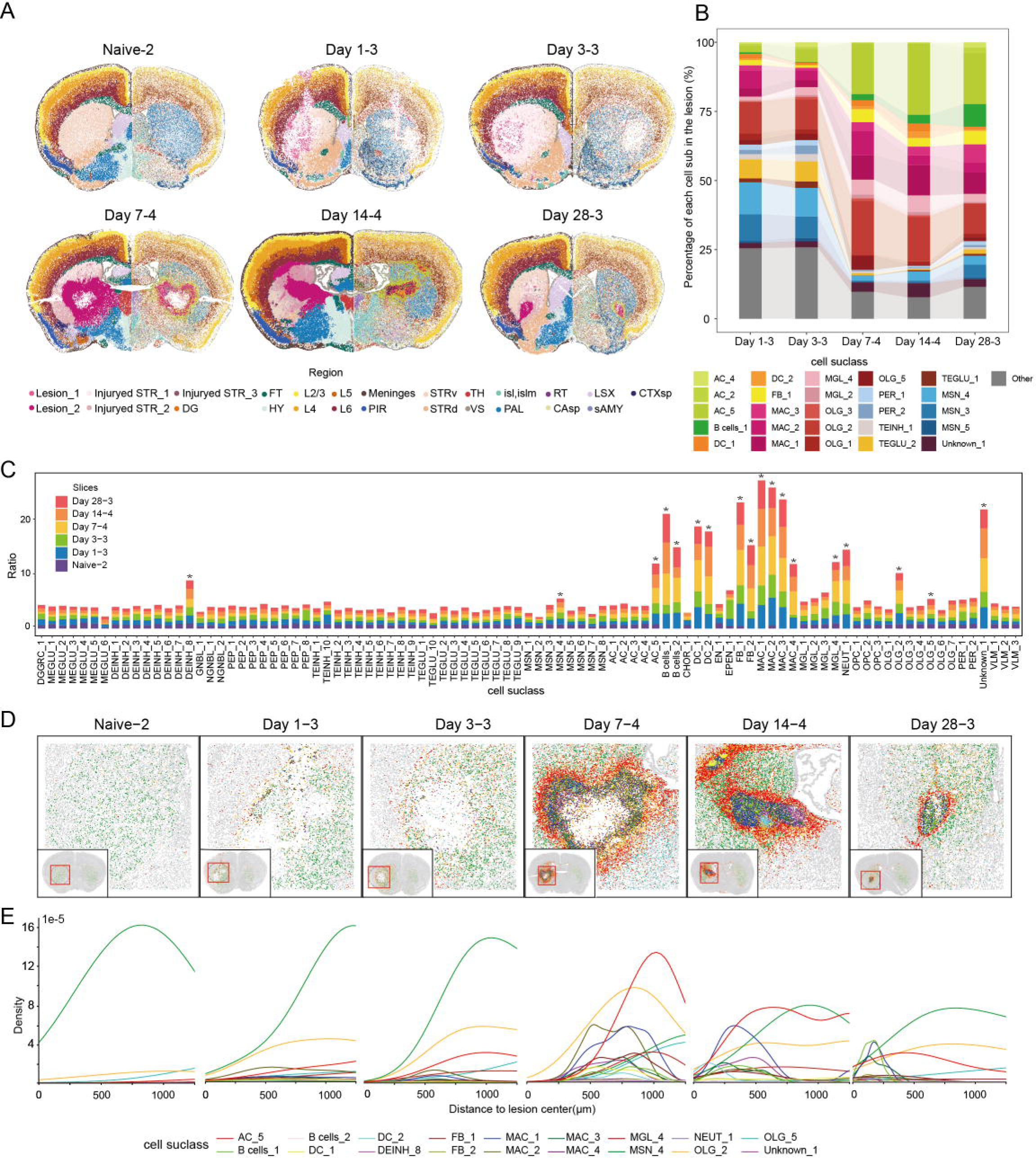
Spatiotemporal distribution of injury-associated cell subclasses. (A) The alignment between molecularly defined tissue regions (left, colored based on molecular tissue region identities) and cell subclasses assignment (right, colored based on sub-celltype assignment). Each dot represents a cell. The anatomical definition of the right hemisphere comes from the Allen Brain Atlas. The abbreviation of the tissue region is consistent with the Allen Mouse Brain Reference Map. L2/3, cortical layer 2 and cortical layer 3; L4, cortical layer 4; L5, cortical layer 5; L6, cortical layer 6; lesion_1, lesion 1; lesion_2, lesion 2; Injured_STR_1, injured striatum 1; Injured_STR_2, injured striatum 2; Injured_STR_3, injured striatum 3; DG, dentate gyrus; FT, fiber tracts; HY, hypothalamus; PIR, piriform area; STRd, striatum dorsal region; STRv, striatum ventral region; TH, thalamus; VS, ventricular systems; isl,islm, islands of calleja and major island of calleja; PAL, pallidum. RT, reticular nucleus of the thalamus; CAsp, field ammon’s horn, pyramidal layer; LSX, lateral septal complex; sAMY, striatum-like amygdalar nuclei; CTXsp, cortical subplate. (B) Cellular composition of the lesion at different time-points post-ICH. Only cell subclasses with a representation of at least 0.5% in the lesion at two or more time-points are individually labeled. Cell subclasses not meeting this threshold are collectively designated as ‘Other’. (C) The density ratio of different cell subclasses between the left (injured) and right (uninjured) hemispheres at time points. The densities of 18 cell subclasses were significantly higher in the left hemisphere compared to in the right hemisphere. \**P*<0.05 vs. right hemisphere (paired Wilcoxon test) (D) Spatial visualization of injury-associated cell subclasses at different time points within the lesion, with each dot representing an individual cell. The cell color codes are the same as those used in the Figure 2E. Using an interactive website(https://db.cngb.org/stomics/stmich/), one can zoom in and display specific cell subclasses. (E) An illustration of the variation in density of injury-associated cell subclasses in relation to the distance from the lesion center.

We performed statistical analysis to identify the cell subclasses involved in the cellular responses to ICH. To accomplish this, we compared the number of cells per unit area of each cell subclass in the injured and uninjured hemispheres. Our analysis revealed that the population size of 18 out of the 92 cell subclasses was higher in the injured hemisphere compared to the uninjured hemisphere (**Figure 2C**). These 18 cell subclasses were defined as ICH-associated cell subclasses. We then analyzed their spatial distribution (**Figure 2D** and **2E**). In the naïve brain, we see mainly neurons (MSN_4), and some oligodendrocytes (OLG_2,OLG_5) and astrocytes (AC_5). During the first 3 days after ICH, the hematoma and neuronal cell death produced a cavity at the bleeding site. Surviving neurons (MSN_4) and astrocytes (AC_5), oligodendrocytes (OLG_2), microglia (MGL_4), and macrophages (MAC_1, MAC_2, MAC_3, MAC_4) were present around the hematoma. From day 7, the lesion was surrounded by a glial scar consisting mainly of astrocytes (AC_5), with many interspersed oligodendrocytes (OLG_2). Astrocytes were primarily located at the lesion border, while oligodendrocytes were more widely distributed. Macrophages (MAC_1, MAC_2, MAC_3, MAC_4) were the predominant cell type in the lesion core, along with some fibroblasts (FB_1, FB_2). A few neutrophils (NEUT_1), dendritic cells (DC_1, DC_2) and B cells were also detected within the lesion.

### Glia cell activation dynamics and functional diversity in response to ICH

In response to ICH, microglia, astrocytes and oligodendrocytes are known to become activated driving proliferation^1,8,34^. We examined the responses of microglia, astrocytes and oligodendrocytes using signature marker genes (e.g., *Gfap* and *C4b* for astrocytes; *B2m* and *Lyz2* for microglia; *Il33* for oligodendrocytes) identified in previous studies^6,35–37^. As expected, there was an increase in the activation of these glial cells within and around the lesion (**Figure S5A**). The degree and state of activation varied across different time-points; the molecular responses were relatively mild during the first three days and became more pronounced on 7 days after ICH. In addition to the activation observed within the lesion, we also noted activation of astrocytes in the corpus callosum. The glial cell subclasses MGL_4, AC_5, and OLG_2 exhibited higher activation scores more proximal to the bleeding site **(Figure 3A-3F)**. These three glial cell subclasses are strongly associated to ICH, exhibiting higher density in the injured hemisphere **(Figure 2C)**. We therefore refer to these cell subclasses as activated glial cell states. Similar to microglia, all macrophage subclasses, demonstrated increased molecular activation scores (**Figure 3D**).

**Figure 3.**
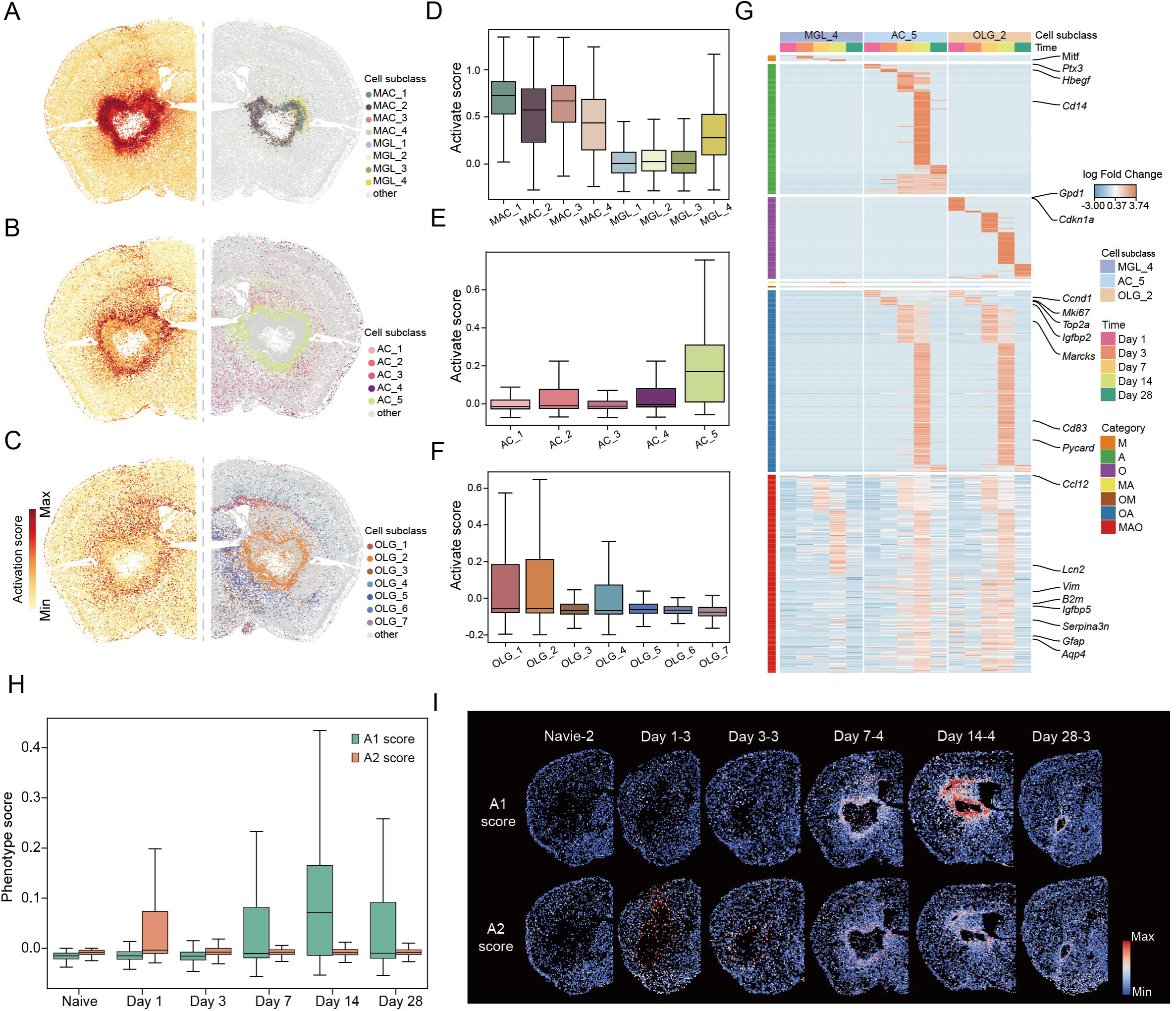
Glial cell activation dynamics and functional diversity. (A) Spatial visualization of activation scores in injured hemispheres on day 7, slice 4 (left) and the subclasses of microglia and macrophages in the injured hemispheres on day 7, slice 4 (right). (B) & (C) Similar to the format in (A), (B) and (C) showcase astrocytes and oligodendrocytes respectively. (D-F) Boxplot representation of activation scores across macrophage and microglia (D), astrocyte (E) and oligodendrocyte (F) subclasses. (G) Heatmap depicting the log-fold change in the expression levels of genes upregulated in the activated glial cell subclasses at the five time-points after ICH compared to the naive group. M: microglia-specific genes; A:astrocyte-specific genes; O: oligodendrocyte-specific genes; MA: genes upregulated in both microglia and astrocytes; OM: genes upregulated in both microglia and oligodendrocytes; OA: genes upregulated in both astrocytes and oligodendrocytes; MAO: genes upregulated in all three glial cell types. (H) A boxplot illustrating the phenotypic scores of A1 and A2 astrocytes within activated astrocyte subclasses (AC_5) at various time intervals: naive, day 1, day 3, day 7, day 14, and day 28. Genes utilized for estimating astrocyte A1 phenotype scores were: *C3*, *H2-T23*, *Serping1*, *H2-D1*, *Gbp2*, *Fkbp5*, *Psmb8*, *Srgn*, and *Amigo2*. Genes utilized for estimating astrocytes’ A2 phenotype scores were: *Clcf1*, *Tgm1*, *Ptx3*, *S100a10*, *Sphk1*, *Cd109*, *Ptgs2*, *Emp1*, *Slc10a6*, *Tm4sf1*, *B3gnt5*, *Cd14*, and *Stat3*. (I) Spatial visualization of A1/A2 astrocyte scores in injured hemisphere across selected brain sections, including naive-2, day 1-3, day 3-3, day 7-4, day 14-4, and day 28-3.

To identify the expression dynamics of genes upregulated in the activated glial cell subclasses, we performed differential gene expression analysis at different time-points after ICH and compared these to the molecular signatures of the corresponding naive cell states. Our study has revealed distinct cell type specific temporal molecular trajectories in gene expression (**Figure 3G**, **see methods**). For example, at 14 days post-ICH, microglia exhibited upregulation of *Mitf*, coding for a transcription factor associated with enhanced phagocytosis^38^. Astrocytes showed upregulation of *Ptx3* and *Hbegf*, particularly in the early stages following ICH (**Figure 3G and S5B**), the protein products of which were shown to support blood-brain barrier integrity during the acute phase of stroke^39^, and play a role in astrocyte morphology, proliferation and differentiation^40^, respectively. In oligodendrocytes, the expression of *Gpd1* and *Cdkn1a* was elevated, particularly on the first day.

We found numerous genes that are co-activated in different cell types, which is an indication of some potential functional overlap between these cells **(Figure 3G and S5C)**. Among these genes, *Ccl12*, a well-known chemokine ligand, was highly expressed in all three glial-cell subclasses clusters on the first day after ICH. Similarly, *Lcn2* and *Aqp4* were also upregulated in all three glial-cell clusters, which may be associated with increased iron ions and edema in the lesion^41,42^. *Serpina3n,* that is associated with neuroinflammation and demyelination^43,44^, showed increased expression levels, particularly in astrocytes and oligodendrocytes after ICH (**Figure S6A**). *Vim* and *Gfap*, genes that are known to be upregulated in reactive astrocytes^45^, were also found to be upregulated in microglia and oligodendrocytes after ICH (**Figure S6A**). Astrocytes and oligodendrocytes upregulated some critical genes involved in cell proliferation (*Mki67, Ccnd1, Top2a*) in the early stages of ICH (**Figure 3G and S6B**). *Marcks*, a pivotal gene for myelin repair^46,47^, showed high expression in oligodendrocytes and astrocytes on day 7 post-ICH **(Figure 3G)**. On day 14 after ICH, genes related to the NF[κB pathway (*Cd14*, *Cd83*, *Bcl3*, *Bcl2a1a*, *Pycard*)^48^ were highly upregulated, in astrocytes (**Figure 3G**).

Our in-depth analysis of astrocytes revealed that these cells have distinct functional adaptations at various stages following ICH. Increased expression of *Igfbp2* was identified in activated astrocytes on day 3, while *Igfbp5* levels were upregulated on day 7 (**Figure S7A and S7B**). On day 3, astrocytes displayed an increased expression of cell cycle-related genes (**Figure 3G**), supporting previous reports that *Igfbp2* enhances proliferation^49^. Our findings also revealed increased expression of C3 in astrocytes on day 14 (**Figure S7A and S7B**). Previous work proposed classification of astrocytes based on their gene expression profile including *C3* as a driver, into neurotoxic (A1) and neuroprotective (A2)^6,50^, even though our recent consensus review strongly advised against using this oversimplifying classification^51^. Using an array of markers to assess A1-like and A2-like phenotypes in the activated astrocytes(AC_5)^52–56^, we found that astrocytes displayed A2-like reparative phenotype mainly on the first day after ICH. In contrast, on day 14 after ICH, astrocytes exhibited pro-inflammatory characteristics, indicative of the A1-like phenotype, in particular in the perilesional regions(**Figure 3H and 3I**). These findings highlight the dynamic shifts in astrocyte functions from an initially neuroprotective to a more neurotoxic at the later phenotype stages after ICH.

### Temporal characterization of molecular changes in the lesion

Over a time span of several weeks to months after ICH, the blood clot is cleared and the damaged tissue partially repaired. To identify gene programs involved in repair and functional recovery, we utilized the Hotspot algorithm^57^, a tool that assesses non-random variations. This tool has enabled us to recognize gene groups that work in synergy to perform specific functions, commonly called modules.

Using hotspot analysis, we have identified 55 gene modules in the lesion and the injured striatum (**Table S5**). To better understand the possible biological functions of these gene modules, we conducted Gene Ontology (GO) analyses (**Table S5**) and identified some gene modules in both naive and ICH groups linked to normal brain functions. For example, GO analysis for module 4, which is located in the striatum, has shown an association with the regulation of the transport of calcium and other ions (**Figure S8A and S8B**).

We manually annotated 25 gene modules associated with the lesion (**Figure 4A and S8C, Table S5**) and the enriched GO terms corresponding to these gene modules are illustrated in **Figure 4B** and **S8D**. We identified specific modules ubiquitous across all five time points after ICH, such as module 10, which contains a set of chemokine genes involved in the recruitment and migration of immune cells. We also noted temporal patterns in some modules. The primary GO terms related to wound healing (module 42) were observed on day 1 after ICH. Module 42 includes a cluster of acute-phase genes that exhibit early changes upon cellular activation and gradually return to baseline levels **(Figure S9A)**. AC_5 exhibited higher expression of module 42 genes, indicating that astrocytes were among the earliest cell types to respond to injury **(Figure S9B)**. Module 37, enriched in terms related to the cell cycle, exhibited elevated scores on days 3 and 7, implying that the time between the day 3 and day 7 represents cell proliferation at this period after ICH (**Figure S9C**). However, it is noteworthy that, on day 3, in module 37, MSN also expressed cell cycle-associated genes, likely induced by neuronal cell death pathways rather than mitosis^58–61^ **(Figure S9D)**. From day 7 post ICH, we observed enrichment in genes associated with phagocytosis (module 2) and granulocyte differentiation regulation (module 44). Adaptive immune responses were observed on days 14 and 28, involving the T cell receptor signaling pathway (module 12), antigen processing and presentation (module 5), and production of immunoglobulin (module 17, module 20). Module 9 and module 39, both associated with the extracellular matrix, exhibited distinct temporal patterns. Module 9, containing genes involved in collagen fibril organization, was more prominent expressed on day 7 post ICH, while module 39, associated with metallopeptidase activity, was more pronounced on day 28. This finding suggests that the lesion initially undergoes collagen fibril organization, followed by subsequent degradation, mediated by metallopeptidase activity.

**Figure 4.**
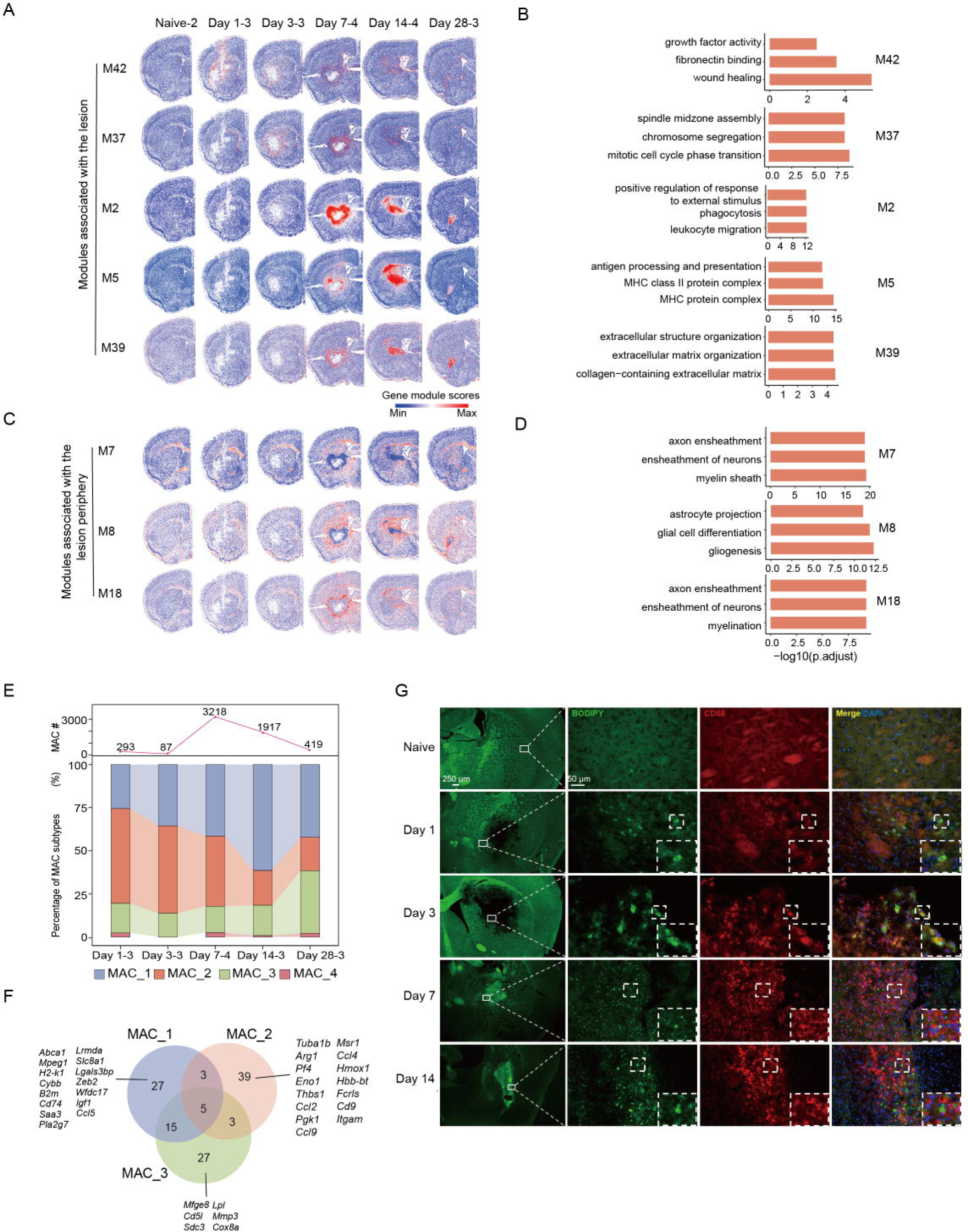
Temporal characterization of molecular changes in lesions. (A) & (C) Spatial visualization of gene modules associated with the lesion, colored according to the module scores. (A) Gene modules within lesions and (C) Gene modules at the lesion periphery. For additional module information, refer to Figure S8A and 8C. Key: ’M’ denotes modules. (B) & (D) Gene ontology (GO) enrichment analysis of gene modules from (A) & (C). X-axis: –log10(p-adjust) score; Y-axis: GO terms. Only top 3 terms (p-adjust < 0.01) are listed for each gene modules. (E) Stacked bar chart with an overlaid line graph depicting the dynamic distribution of macrophage subpopulations over various time-points. The bars represent the percentage of each macrophage subclass within the total population at a given time, while the line graph indicates the total macrophage count, with specific values labeled at key time-points. (F) Venn diagram illustrating the overlap of the top 50 differentially expressed genes in the macrophage subclasses MAC_1 (blue), MAC_2 (red), and MAC_3 (green). Representative genes are enumerated within each section to represent the distinctive and shared genes. Numerical values indicate the count of unique or overlapping genes for each respective subpopulation. (G) Immunofluorescence staining of lipids and macrophages from naive, day 1, day 3, day 7 and day 14 after ICH. Macrophages were targeted using an antibody against CD68. Lipids were marked with BODIPY.

We identified three gene modules associated specifically with the lesion periphery (**Figure 4C**). Gene modules 7 and 8 contain genes enriched in GO terms related to the myelin sheath, ensheathment of neurons, and gliogenesis (**Figure 4D**).

Furthermore, module 18 comprises of a set of genes related to neurogenesis and axon myelination regulation (**Figure 4D**), such as *Sox10*, which is expressed in myelin-forming oligodendrocytes^62^, and *Plxnb3,* the protein product of which promotes neurite outgrowth^63^. This module was detected around the lesion and was most prominent on days 7 and 14. These observations reveal the time window for intitiation of remyelination and axon regeneration in the perilesional region following ICH.

Histochemical visualization of myelin sheath showed cavities and demyelination in the lesion area from day 1 to day 3. Cells gradually started populating these cavities on day 7 and we observed at the lesion periphery ongoing remyelination that was characterized by a shallow gradient, finer and shorter myelin fibers, and an arrangement either in a crossing pattern or along the fiber center. Remyelination became more complete, extending towards the lesion edge, in the period from days 14 to 28 after ICH. These findings provide insight in the temporal dynamics of demyelination and remyelination in the lesion periphery following ICH (**Figure S9E**). Macrophages (MAC) are essential for hematoma clearance and tissue remodeling. In an attempt to further characterize their involvement, heterogeneity and dynamics, we examined all four subclasses of macrophages in a spatiotemporal manner **(Figure 4E and S9F)**. MAC_2 cells were most abundant in the period from days 1 to 3 after ICH, while the MAC_1 and MAC 3 cells constituted the largest MAC subtype on day 14 and 28, respectively. The number of MAC_4 cells was relatively low at all investigated time points. MAC_2 molecular signature was enrichment in pro-inflammatory genes, such as *Ccl4, Ccl9, Ccl2,* and *Pf4*, whereas MAC_3 cells are characterized by the expression of *Mmp3*, *Mfge8*, and *Cd5l* **(Figure 4F)**. Mmp3 is known to participate in the degradation of multiple ECM components^64^ and Mfge8 is involved in clearing of apoptotic cells and regulation of inflammatory processes^65–67^. These findings suggest that MAC_3 represents a subclass of macrophages responsible for clearing of extracellular matrix excess and death cells. The MAC_1 populations was characterized by phagocytic vesicle-related genes, MHC protein complex genes, and lipoprotein particle complex genes such as *Abca1, H2-K1, Mpeg1, Cybb, B2m, H2-D1, Cd74, Saa3, Pla2g7,* and *Pltp* **(Figure 4F)**. This molecular signatures suggest that MAC_1 represent a subset of macrophages overloaded with lipids, similar to foam cells. To confirm the presence of foam cells in the lesion, we co-stained brain sections with BODIPY fluorescent dye to label lipids and anti-CD68 antibody to visualize macrophages (**Figure 4G**). In the control tissue, there was almost no detection of lipids in macrophages. On days 1 and 3 post-ICH, there was a slight increase in the colocalization of these markers within the lesion. From day 7 to day 14 after ICH, there was a significant accumulation of lipids in the lesion area, with macrophages containing large numbers of lipid droplets. These findings highlight the dynamic shifts in macrophage phenotypes after ICH and underscore their diverse roles and adaptations in the inflammatory response, ECM remodeling, and lipid metabolism within the lesion area.

### Spatial variations in gene expression within the lesion

Next, we explored whether there were any spatial patterns in gene expression within the lesion. On day 7 post-ICH, the lesion formed a closed spatial entity with peak expression levels of scavenger receptors (**Figure S10A**). Usig a generalized linear model, we therefore analyzed the relationship between gene expression patterns and their proximity to the lesion center at this time point(**Figure 5A**).

**Figure 5.**
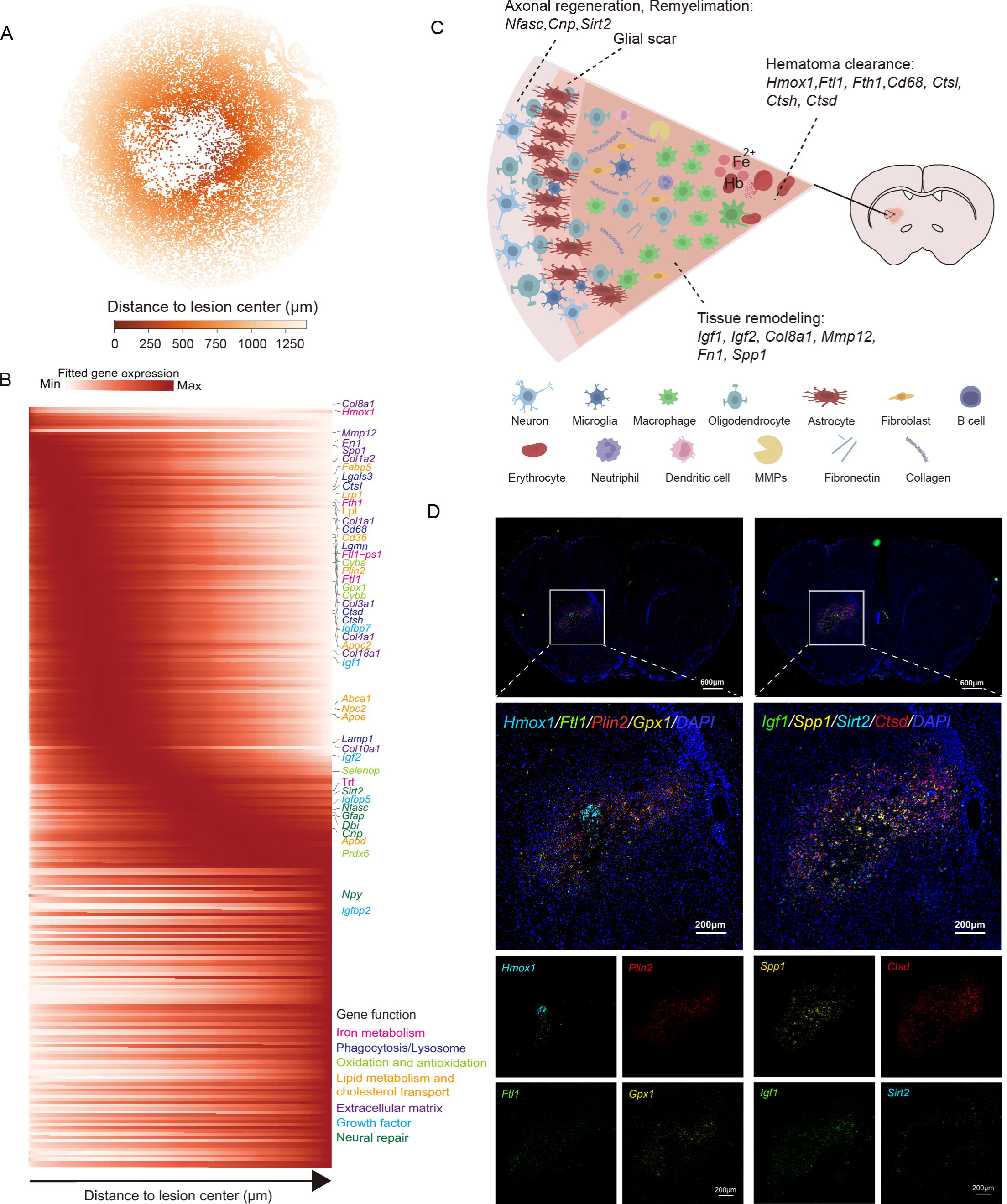
Spatial variations in gene expression within the lesion. (A) Spatial distribution of cells around the lesion site on day 7 post-ICH. The lesion is represented as a circular region with a 1400 µm radius. Each point marks the position of a cell, with the color gradient reflecting the cell’s distance from the center of the lesion. (B) Heatmap of gene expression gradients on post-ICH day 7. The heatmap displays variations in expression levels as a function of the distance from the lesion center. The text color of genes is depicted according to their potential functional categories. (C) Illustrative view of lesion components. (D) Fluorescence In Situ Hybridization (FISH) results for specific genes (*Hmox1, Plin2, Spp1, Ctsd, Ftl1, Gpx1, Igf1* and *Sirt2*) in the lesion on day 7 post-ICH.

Several genes showed a spatial gradient from the lesion core on day 7 after ICH. For example, *Hmox1*, coding for an enzyme crucial to the breakdown of heme, was expressed near the lesion center, consistent with our previous report^68^. Moreover, genes involved in phagocytosis, such as *Cd68, Lrp1, Cd36, Ctsl, Ctsh, Ctsd, Lamp1, Lgmn,* and *Lgals3,* were also enriched near the lesion center **(Figure 5B and S10B**). *Ftl1* and *Fth,* the protein products of which facilitate the storage of excess ferric irons following hemoglobin/heme breakdown, were mainly expressed by immune cells close to the lesion area **(Figure S10C**). The gene products of antioxidant-related genes like *Gpx1*, *Selenop* were mainly expressed in macrophages and other immune cells near the lesion center and *Prdx6* was primarily expressed in astrocytes closer to the lesion border (**Figure 5B and S9D**). These increases in antioxidant gene expression could neutralize the excess ROS from active inflammation and Fenton reaction after ICH^69–71^. Lipid and cholesterol metabolism genes, including *Apoe, Lpl, Abca1, Plin2, Apoc2, Npc2,* and *Fabp5,* were also upregulated within the lesion center as well as components of the extracellular matrix, such as *Col8a1, Mmp12, Fn1, Spp1, Col1a2, Col1a1, Col3a1, Col4a1, Col10a1, Col18a1,* and together with growth factor members coding genes including *Igf1, Igf2, Igfbp5, Igfbp7, Igfbp2* (**Figure 5B, S10F and S10G)**. In the lesion periphery, we observed the expression of *Nfasc, Cnp*, and *Sirt*2, associated with neural repair **(Figure 5B and S10H)**. The cell adhesion protein

Nfasc is involved in maintaining the axon initial segment, impulse conduction, and the assembly of the nodes of Ranvier^72–74^. Its expression suggests the potential for promoting the formation of initial segments of axons and initiating action potentials around the lesion. Additionally, both *Cnp* and *Sirt2* demonstrated predominant expressions in oligodendrocytes and OPC. Sirt2 has been proven to enhance remyelination^75^, while Cnp can mediate process outgrowth in oligodendrocytes^76^. These findings suggest that *Nfasc, Cnp, and Sirt2* could participate in repairing and renewing myelin sheaths near the lesion. Finally, based on our data, we constructed an illustration summarizing the pathophysiological processes that occur 7 days post-ICH(**Figure 5C**). The discovery potential of our spatial transcriptome capture was confirmed by fluorescent *in situ* hybridization (FISH) of selected representative genes **(Figure 5D)**.

### Impact of ICH on different brain regions

To evaluate the impact of ICH on brain regions away from the lesion, we employed two distinct comparative analyses (**Figure 6A)**. The initial analysis juxtaposed ICH-impacted brains with naive controls, thereby elucidating the differences between affected and unaffected cerebral landscapes. Subsequently, the second analysis contrasted the ipsilateral (injured) with the contralateral (uninjured) hemispheres within the same cerebral section to identify the asymmetric effects of ICH on brain functionality.

**Figure 6.**
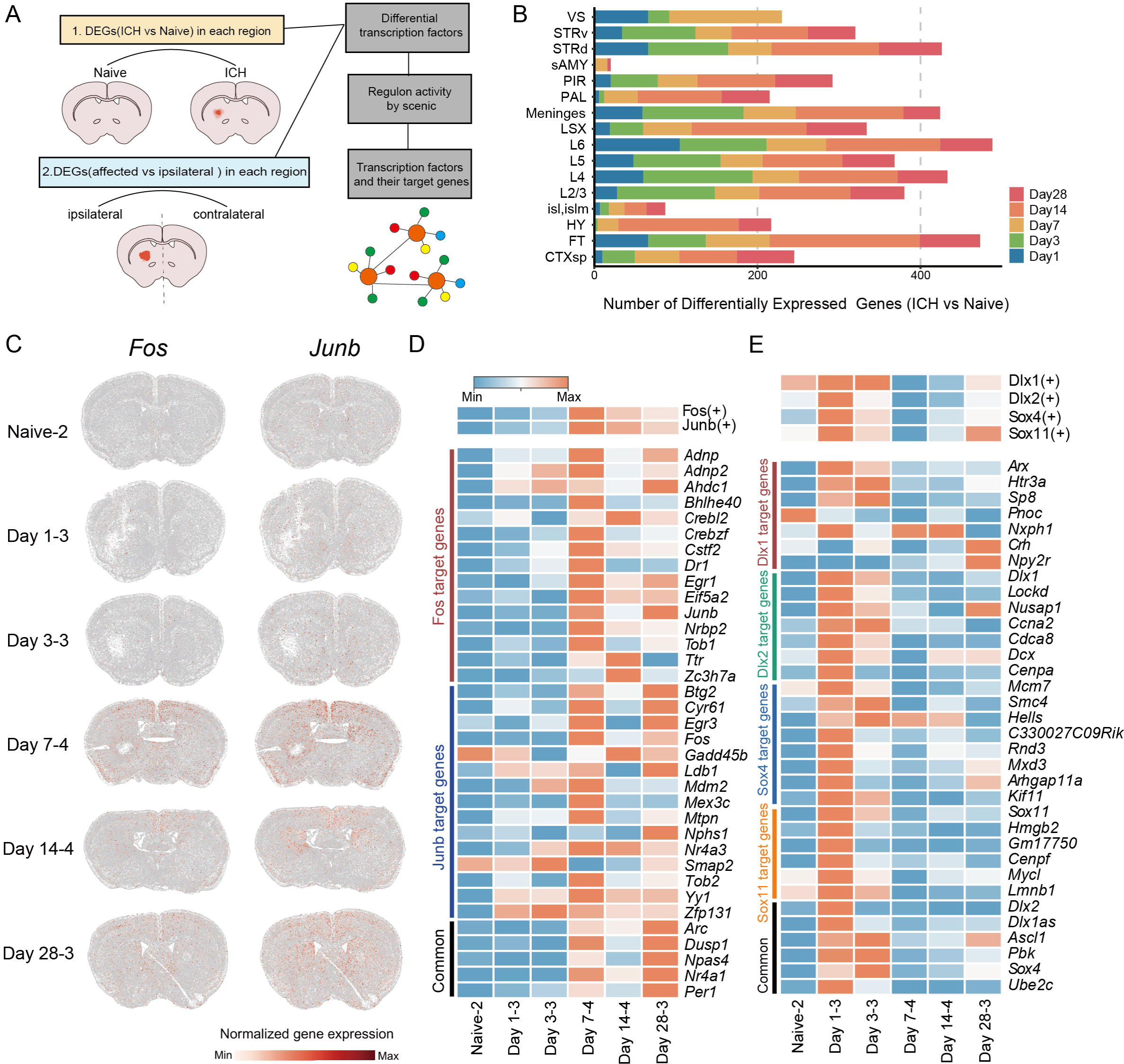
Molecular changes in different brain regions after intracerebral hemorrhage. (A) Schematic representation of the comparative analyses. (B) The number of differentially expressed genes in various brain regions at different time points following ICH compared to the naïve group. Significant genes were identified based on specific criteria: logfoldchange≥1, scores > 5, and p-adjust< 0.001. (C) Spatial distribution of *Fos* and *Junb*. (D) Heatmap illustrating the transcriptional regulatory activity of regulons (Fos and Junb), and the column-scaled mean expression of target genes for regulons (Fos and Junb) in the cortex of the naive group and at each time point after ICH. (E) The regulon activity of neural stem cell activation-related regulators, and the column-scaled mean expression of target genes for regulons (*Dlx1*, *Dlx2*, *Sox4*, *Sox11*) in the VS region of the naive group and at each time point after ICH.

We found that the variation in gene expression was more pronounced between ICH-affected brains and naive controls compared to the variation between ipsilateral and contralateral regions **(Figure 6B and Figure S11A, Table S6)**. This variation in gene expression can be attributed partly to the differences in spatial localization of the affected areas, and partly to the pervasive, global influence that ICH exerts on the brain’s molecular landscape. For further analyses, we focused on regions common to all tissue sections, such as the cerebral cortex, ventricular systems (VS), and fiber tracts (FT).

Following the analysis of gene expression across different cortical layers in ICH-affected versus naive brains, we found that the main effects were linked to time after ICH rather than cortical layer function or structure. ICH led to similar molecular changes across all cortical layers, highlighting a common molecular response to ICH (**Figure S11B**). After day 7 post-ICH, AP-1 family transcription factors, including *Junb* and *Fos*, were upregulated across cortical layers (**Figure 6C**). To further elucidate their regulatory functions, we utilized Single-cell Regulatory Network Inference and Clustering (SCENIC)^77^, enabling the computation of transcription factor activity while mitigating issues such as dropout events and technical variability. This enhanced the robustness and reliability of our analyses. The regulatory activity of Junb/Fos aligned with the observed increased expression of immediate early genes (IEGs) such as *Arc*, *Egr1*, *Nr4a1*, and *Npas4* (**Figure 6D**), that play pivotal roles in brain plasticity and memory formation^78–82^. Given that chronic activation of IEGs has been linked to synaptic loss and cognitive impairment^83^, our findings also point to a possible mechanistic pathway to long-term neurological deficits induced by ICH.

We performed GO enrichment analysis on differentially expressed genes across cortical layers after ICH (**Table S7**). Across cortical layers at the same time-point, we identified similar GO terms. In contrast, significant variations in GO terms were observed across different times within the same cortical layers. For example, similar biological processes were increased at layer 4 on days 1 and 3, including regulation of neuronal death, acute inflammatory response, and positive regulation of phagocytosis. Starting from day 7, layer 4 exhibited enrichment in genes regulating lipid transport and regulation of neuronal synaptic plasticity **(Figure S11C)**.

In the VS region on days 1 and 3 post-ICH we noted elevated expression of transcription factors/regulons associated with NGNBL activation in naïve control animals^84^ (**Figure 6E and Figure S11D**). The target genes of these regulons primarily encompassed GO terms related to mitosis, cell cycle, neurodevelopment and GABAergic interneuron differentiation (**Table S7**). Activated NGNBL were differentiating into new neurons, indicated by an upsurge in the number of cells expressing *Dcx*, a marker of immature neurons^85^(**Figure S11E**, **Table S6**).

The comparative analysis between the ipsilateral and contralateral hemisphere revealed a smaller number of differentially expressed genes (**Figure S11A, Table S6**). Temporally, the first day post-injury showed a higher number of differentially expressed genes, with GO enrichment primarily related to neuronal death and the regulation of immune processes. Over time, the gene expression profiles of the two brain hemispheres gradually converged. For instance, by day 28 after ICH, there were no differentially expressed genes in the ipsilateral and contralateral layer 4. In contrast, the corpus callosum, a white matter fiber tract connecting the brain’s two hemispheres, showed asymmetrical expression of genes associated with cell migration and immune response (**Table S7**), such as *C4b*, *Serpina3n*, *Klk6* and *Gfap* (**Figure S11F**).

## Discussion

We employed Stereo-seq to conduct a spatial transcriptomic analysis of mouse brain at critical time points after ICH. To our knowledge, this is the first spatiotemporal map of ICH at single-cell resolution. This study has identified and investigated 23 major cell types and 92 derived subclasses, advancing the understanding of the cellular landscape in ICH-induced brain injury. This detailed analysis provides crucial insights into ICH-induced injury mechanisms and genes involved that will assist the development of novel therapeutic approaches targeting specific pathological or regenerative processes.

Following ICH, we fully characterized the lesion and its surrounding, identified a wide range of cells and there adaptations. Our quantitative analysis revealed an augmented presence of 18 cell subclasses in the injured hemisphere, contributing to the progression and recovery of the lesion. Oligodendrocytes significantly repopulate the areas around the lesion after 7 days and persist over time, which relates to the repair of myelin sheaths around the lesion. The near complete absence of neurons in the lesion area from day 7 to day 28 underscores the challenges in neuronal survival and limited ability for neuronal netoworks to regeneration following ICH.

Astrocytes, microglia and oligodendrocytes display activation signatures, which exhibits a pattern of acute induction followed by subsequent attenuation. Astrocytes form dense scar tissue in the acute phase of injury, restricting the lesion area and isolating the injured tissue from its healthy surroundings. However, this dense scar may hinder regeneration as the hematoma is cleared by phagocytic cells. From a cellular and molecular perspective, we discovered a dual role of astrocytes, which indicates molecular transformation. Astrocytes exhibit protective properties in the early stages after ICH, but in the later stages, they attain molecular signatures that can drive neurotoxicity. This process is also accompanied by a transition of astrocytes from a proliferative to a non-proliferative state, as observed in this study and recently reported by others^86^. Notably, the Insulin-like Growth Factor Binding Protein (IGFBP) family members appear to play a significant role in this transition. In the first 3 days following ICH, IGFBP2 promotes the proliferation of astrocytes. However, after day 7, astrocytes exhibit a non-proliferative phenotype, which may be related to the expression of IGFBP5. IGFBP5 has been reported to inhibit the proliferation of astrocytes^49^. Pharmacological intervention preventing astrocyte transition from protective to neurotoxic could enhance the subsequent regeneration.

We identified a series of complex processes that entail early peripheral immune cell recruitment and proliferation of various cell types in the ICH lesion. These processes are followed by hematoma phagocytosis, extracellular matrix production, and changes in the immune response between 14-28 days. The later phase is characterized by antigen presentation, immunoglobulin production, and B cell activation as well as lipid accumulation and the formation of foam cells. Given that foam cells can contribute to a chronic inflammatory environment^87^, targeting foam cells or preventing lipid overload could help brain repair after ICH. Notably, recent research has demonstrated that using liver X receptor agonists to promote cholesterol efflux and macrophage recirculation can reduce brain injury and promote tissue regeneration following ICH^88^. This injury microenvironment bears similarities to atherosclerosis, where macrophages engulf oxidized low-density lipoprotein (oxLDL) and transform into foam cells. OxLDL has been shown to induce B cell activation and the production of natural antibodies that bind to oxLDL^89,90^. Following ICH, the breakdown of damaged myelin sheaths generates a considerable amount of polyunsaturated fatty acids, free cholesterol, cholesterol esters, and phospholipids. These lipids can be loaded onto low-density lipoprotein (LDL) particles. The inflammatory environment facilitates various lipid peroxidation pathways, increasing the likelihood of LDL oxidation and subsequently inducing B cell activation.

Our study shows increased expression at the lesion border of several genes coding for proteins that promote axonal regeneration and myelin repair, including *Nfasc*^72–74^, *Sirt2*^75^, *Cnp*^76^, *Sox10*^62^, and *Plxbn3*^63^. Furthermore, we observed the activation of neural stem cells in the subventricular zone (SVZ), accompanied by an increase in the generation of new neurons, as indicated by expression of *Dcx*. Previous studies reported that injuries such as stroke increase precursor proliferation in the SVZ and induce ectopic migration of SVZ-derived cells to the injury site^91–94^. This finding indicate that the brain possesses an inherent capacity for self-repair following injuries that cause extensive neuronal death. Although it is unclear whether the newly generated neurons can integrate into existing neural circuits, we observed the activation of AP-1/IEGs on post-ICH days 7, 14, and 28. The activation of IEGs regulates synaptic plasticity and promotes neural circuit remodeling, implying that extensive neural remodeling may occur after ICH^95–99^. Nevertheless, chronic AP-1 activation during aging can promote human tau pathology and degeneration^100^.

## Acknowledgments

M.W. is supported by grants from the National Key R&D Program of China (2022YFC3400300).This work was supported partially by the National Natural Science Foundation of China (No. 82371339 to Jian W.). L.L. is supported by the Shenzhen Basic Research Project for Excellent Young Scholars (RCYX20200714114644191) and the National Key R&D Program of China (2022YFC3400405). Project analysis was performed partially on the STOmic Cloud (https://cloud.stomics.tech). M. Pekny is supported by grants from the Swedish Research Council (2020-01148, 2019-00284), Hjärnfonden (FO2021-0082) and ALF (965939), M. Pekna is supported by grants from the Swedish Research Council (2021-01486), Hjärnfonden (2020-0134) and ALF (966011).We thank the National Supercomputing Center in Zhengzhou for its computational resources. This work was also supported by China National GeneBank (CNGB). JY Chen and YF Yang are contribute to draw the picture in the manuscript. This paper is from the Mesoscopic Brain Mapping Consortium.

## Author contributions

Jian W., M.W., Junmin W., and X.C. conceived the study. M.W., Z.T., R.D., Z.J., and S.W. performed experiments. R.X., Z.C., and J.T. analyzed and interpreted the data. T.Y. and J.C. constructed the database. Q.P., X.L., and X.Z. performed the verification experiments. L.L., J.M., Milos P., and Marcela P., interpreted the data, U.W., Y.P., N.M., X.F., C.L., X.X. and C.J. provided feedback. S.C., N.C., M.Z., J.L., and Q.X. drew and composed the images. R.X., Z.C., J.T., Junmin W., Jian W., and X.C. drafted and revised the manuscript. All authors reviewed,edited and approved the manuscript.

## Limitations

It is imperative to acknowledge the limitations of our study, particularly the inconsistency of brain section preparation. Despite our efforts to ensure uniformity in sample preparation, we encountered significant challenges in achieving precise alignment of brain sections, which resulted in minor variations in the spatial and molecular data. It is worth noting that this task is complex, so it is essential to exercise caution when interpreting the data. Nonetheless, our findings are still valuable and contribute to the existing knowledge on this subject.

## Materials and Methods

### Mouse ICH model and tissue collection

#### Animals

A total of 24 C57BL/6 male mice (3-month-old, 22-25g) were procured from SPF (Beijing) Biotechnology Co. Ltd. (SCXK(JING)2019-0010). The mice were housed in a specific pathogen-free (SPF) environment with a temperature range of 23-25□ and relative humidity between 45–55%. They were subjected to a 12 h light and 12 h dark cycle and had free access to food and water ad libitum^23^. The Animal Ethics Committee of Zhengzhou University and BGI-research approved the animal protocol (ZZUIRB 2022-148, BGI-IRB E23025). All animal experiments in this study followed the national and the ARRIVE guidelines (http://www.nc3rs.org.uk/arrive-guidelines). The mice were randomized into groups utilizing a website (www.randomization.com).

#### ICH Mouse Model

In this study, anesthesia was administered to male C57BL/6 mice using isoflurane (70% N_2_O and 30% O_2_; 4% induction, 2% maintenance). The mice were then placed in a stereotaxic instrument, and a 1-1.5 cm incision was made along the sagittal suture. After being filled with paraffin wax, the glass micropipette was attached to the micro-infusion pump, and collagenase VII-S (0.075 U in 0.5 µL sterile saline, Sigma, St. Louis, MO) was injected into the left striatum using dual-arm brain stereo locator (anterior to bregma 0.8 mm, lateral to the midline 2.0 mm, deep to the surface of the skull 3.5 mm) (Stoelting, 51503D, Stoelting Co., Illinois, Wood Dale, USA). The collagenase was infused with a constant rate of 0.1 µL/min using a micro-infusion pump, Stoelting, Wood Dale, IL, USA)^101^. The mouse body temperature was maintained at 37.0 ± 0.5□ throughout the operation with the help of a body temperature maintenance instrument (RWD, Shenzhen, China). The control group included normal C57BL/6 male mice; any mice that died within 24 hours after surgery were excluded from the analysis.

#### Brain lesion volume measurement

Three days after ICH, the mice were anesthetized with 3% isoflurane, followed by saline and 4% paraformaldehyde perfusion through the left ventricle. We removed the brain and took coronal sections across the striatum. These sections were stained with Luxol fast blue to allow myelin dyeing. The lesion volume was measured using ImageJ 1.54f. An investigator who was unaware of the experimental cohort conducted the analysis.

#### Brain edema measurement

Three days after ICH, the mice were euthanized under anesthesia. The brain was then removed and divided along the sagittal fissure into the ipsilateral and contralateral hemispheres and cerebellum. The brain tissue was weighed to determine its wet weight, and it was placed in an oven at 100□ for 72 h to obtain its dry weight. Brain edema was calculated by determining brain water content using the formula: brain water content (%) = (wet weight – dry weight)/wet weight×100%^100,101^.

#### Neurologic deficit score

Per our established protocol, we evaluated mouse neurologic deficit score (NDS) on days 1, 3, 7, 14, and 28 after ICH. The NDS system consisted of six parameters that were used to assess the naive and ICH mice: body symmetry, gait, climbing, circling behavior, front limb symmetry, and compulsory circling. Each parameter was scored on a scale of 0 to 4, with a maximum deficit score of 24. Mice with a score of less than 4 after 1 day of ICH were excluded from this study.

#### Tissue collection

For this study, mice were selected based on their NDS. From each group, one mouse that met the criteria per group was chosen for anesthesia, brain retrieval, and subsequent freezing. The cryopreserved brain tissues were then embedded in blocks and sectioned coronally at a thickness of 10 μm using a Leica CM1950 cryostat. We collected tissue sections containing the hemorrhagic region to ensure uniformity in the sampled area across all groups. Each sample comprised 5 tissue sections spaced 100 μm apart along the anterior-posterior axis. Following cryosection, we promptly applied the sample sections to Stereo-seq chips.

### Stereo-seq library preparation and sequencing

#### Tissue processing and imaging

The tissue sections were processed and imaged based on Chen et al.’s protocol for Stereo-seq library preparation and sequencing^20^. The Stereo-seq capture chip was cleaned with RNase inhibitor (NEB, M0314L), enriched with NF-H_2_O, and then air-dried at room temperature before use. The sections were then attached to the chip surface and incubated at 37°C for 3 minutes. The sections were fixed with methanol and set at -20°C for 40 minutes, after which the Stereo-seq library was prepared. The chip-mounted sections were treated with a fluorescent staining solution containing 0.1 × SSC (Thermo, AM9770), 1/200 nucleic acid dye (Thermo Fisher, Q10212), and 2 U/μl RNase inhibitor for 3 min. After removing the staining solution, chips were cleansed with a Wash Buffer containing 0.1 × SSC and 2 U/μl RNase inhibitor. Before in-situ capture at the FITC channel (objective 10×), imaging was carried out using a Motic Custom PA53 FS6 microscope.

#### In situ reverse transcription

The tissue samples on the chip received treatment with 0.1% pepsin (Sigma, P7000) in a 0.01 M HCl buffer, followed by incubation at 37□ for 6 min to ensure permeabilization. The samples were then washed twice with 0.1 × SSC buffer (Thermo, AM9770), to which 0.05 U/ml RNase inhibitor (NEB, M0314L) was added. The RNA molecules trapped by the DNB on the chip and released from the tissue were reverse transcribed by incubating them overnight at 42□ with the reverse transcription mixture (10 U/ml SuperScript II reverse transcriptase, 1 mM dNTPs, 1 M betaine solution PCR reagent, 7.5 mM MgCl_2_, 5 mM DTT, 2 U/ml RNase inhibitor, 2.5 μM Stereo-seq-TSO, and 1 × first-strand buffer). After reverse transcription, the chip was washed twice with 0.1 × SSC buffer and then incubated in the Tissue Removal buffer at 37□ for 30 min. The chips housing cDNA were collected by treating the chip with Exonuclease I (NEB, M0293L) for 1 h at 37[. The residual cDNA was then collected by a final rinse of the chip with 0.1× SSC buffer. The wash buffer and the production of Exonuclease I were combined and purified by 0.8 × VAHTS DNA clean beads.

#### Amplification

After obtaining the resultant cDNAs, we used the KAPA HiFi Hotstart Ready Mix (Roche, KK2602) and a 0.8 μM cDNA-PCR primer to amplify them. PCR reactions were incubated at 95□ for 5 min, followed by 15 cycles at 98□ for 20 sec, 58□ for 20 sec, and 72□ for 3 min. Finally, we completed the process with a final incubation at 72□ for 5 min. After the amplification process, we purified the cDNA using 1 × VAHTS DNA clean beads.

#### Library Construction and Sequencing

Before starting the library construction and sequencing process, we measured the PCR product concentration using the Qubit dsDNA Assay Kit (Thermo, Q32854). We then fragmented 20 ng of cDNA using our in-house Tn5 transposase at 55□ for 10 min and stopped the reaction with 0.02% SDS. After gently mixing the reaction product at 37□ for 5 min post-fragmentation, we added 25 μL of fragmentation product with 20 μL of 5 × KAPA HiFi Hotstart Ready Mix, 3 μL of 10 μM Stereo-seq-Library-F primer, and 3 μL of 10 μM Stereo-seq-Library-R primer and 49 μL of nuclease-free H_2_O. The reaction was conducted with an initial cycle at 95□ for 5 min, followed by 13 cycles of 98□ for 20 sec, 58□ for 20 sec, and 72□ for 30 sec. We then ended the reaction with a final cycle of 72□ for 5 min. Finally, we purified the amplified PCR products using VAHTS DNA clean beads, generated DNB and sequenced them on the MGI DNBSEQ-Tx sequencer.

### Stereo-seq data preprocessing

#### Quality control and alignment of sequencing data

The raw data processing steps were performed using the protocol described previously^20^. We used the MGI DNBSEQ-Tx sequencer to produce Fastq files. The CID and MID sequences can be in read 1 (CID: 1-25 bp, MID: 26-35 bp), while read 2 contains the cDNA sequences. The CID sequences in the first read were initially mapped to the pre-designed coordinates of the in situ captured chip obtained from the primary sequencing round. A single base mismatch was allowed to correct sequencing and PCR errors. Reads with MID containing N bases or more than two bases with a quality score below 10 were removed. Subsequently, the CID and MID of each read were added to the read header. The remaining reads were then aligned to the mm10 reference genome using STAR^103^. Reads with a MAPQ > 10 were annotated and counted for the respective genes. UMIs that shared the same CID and gene locus were combined, allowing a single mismatch to correct sequencing and PCR errors. This information was used to generate the expression profile matrix inclusive of CID.

#### Cell segmentation with nucleic acid staining

We used nucleic acid staining from the same section to segment cells. We projected the staining image onto the Stereo-seq chips and created a spatial density matrix using the total UMI in each DNB spot. This matrix was transformed into an image where each pixel represented a single DNB. We determined each pixel’s grayscale by the total UMI present in the DNB spot. We manually aligned the DNB image with the nucleic acid staining to analyze cell segmentation. Then, we carried out cell segmentation analysis using Cellpose 2.0, with the default model^104^, and extended the cellular nuclei to encompass the cytoplasmic region. Ultimately, the transcripts within a single cell comprised the identified nuclear area and its surrounding 6 μm area.

To assess whether our cell segmentation method delivered single-cell resolution, we evaluated the spatial distribution of nucleus-localized transcripts (*Malat1* and *Neat1*) and cytoplasm-enriched mitochondrial transcripts within individual cells. We identified the central point of each cell and measured the distance between this central point and the nucleus-localized and cytoplasm-enriched mitochondrial transcripts on each DNB. We then computed the distribution of distances from their respective cellular centers for nucleus-localized transcripts and cytoplasm-enriched mitochondrial transcripts across a single brain tissue slice.

### Stereo-seq data analysis

#### Unsupervised clustering and annotation of the segmented cells

Cells containing fewer than 150 detected genes were removed from the dataset. Then, a semi-supervised surgery pipeline was used to preliminarily annotate the remaining cells using a deep learning framework that learned from labeled datasets and annotated the unlabeled datasets. We used Shi et al.’s datasets to construct our model and annotated our datasets accordingly^105^. We removed the batch effect using the scVI algorithm and clustered the cells across all slices using the Leiden algorithm^26,106,107^. We confirmed the cell type identities by validating the resulting clusters using preliminary annotation results and classical markers. Each primary cell type was processed separately using scVI for batch effect removal and the Leiden algorithm for clustering to identify cell subclasses. Marker genes for each subclass were identified by differential gene expression analysis between each subclass and the sampled subsets of all cells.

#### Inferring of signaling pathway activities

We estimated signaling pathway activities for each cell, using the model matrix from PROGENy^108^. We input the normalized data and obtained resources for PROGENy from converted human homologous genes. We fit a linear model for each cell to predict observed gene expression based on the weights of all Pathway-Gene interactions. After fitting the model, we obtained scores from the t-values of the obtained slopes.

#### Spatial domain analysis

We modified the method proposed by Shi et al. and He et al^28,33^. Instead of using the high-dimensional gene expression matrix, we used the scVI method to generate a feature matrix that minimizes the effects of batch differences. Then, we used the feature matrix to generate a smoothed feature vector for each cell by concatenating the vectors of the *k* nearest spatial neighbors, including the cell itself. After that, we combined all the cells’ spatially smoothed feature matrices into one dataset and subjected them to Principal Component Analysis (PCA). We used the Leiden algorithm to cluster the spatial domain^107^. Additionally, we annotated the spatial domain based on the anatomical structures of the Allen Mouse Brain(http://atlas.brain-map.org).

#### Differential expression analysis

To dissect the transcriptomic changes associated with glial activation after ICH, we performed differential expression (DE) analysis on distinct cell populations. At each post-ICH timepoint, activated astrocytes (AC_5), microglia (MGL_4), and oligodendrocytes (OLG_2) were compared to their respective naive states. Datasets exceeding 10,000 observations were randomly subsampled to ensure consistent group sizes and statistical robustness. We employed Scanpy’s rank_genes_groups function to identify DE genes, using stringent filtering criteria: absolute log2 fold change (FC) ≥ 0.64 (equivalent to FC > 1.25), score > 10, and adjusted p-value < 0.0001.

For regional comparisons, we again employed Scanpy’s rank_genes_groups function, but applied slightly relaxed thresholds (logFC ≥ 1, score > 5, and adjusted p-value < 0.001). Two distinct analyses were conducted:

1. ICH vs. Naive: At each post-ICH timepoint, each brain region in the ICH-affected brain was compared to its corresponding region in the naive brain. This identified DE genes associated with ICH pathology in specific brain regions.
2. Ipsilateral vs. Contralateral: Within each post-ICH timepoint, each brain region was further compared between the ipsilateral (injury-adjacent) and contralateral (opposite) hemispheres. This revealed DE genes reflecting local versus non-local responses to ICH injury.

Further gene ontology (GO) enrichment analysis is conducted using the "clusterProfiler" R package^109^. EnrichGO function was applied to each DE gene list with the parameters: pvalueCutoff = 0.05, qvalueCutoff = 0.05, pAdjustMethod = "BH", ont = "ALL".

#### Gene score and cell cycle analysis

Glial activation scores and astrocyte A1/A2 phenotype scores were computed from normalized gene expression data. Leveraging Scanpy’s score_genes function, scores were calculated by summing expression values of specific gene sets, followed by subtracting the mean expression of a randomly chosen background gene set. For astrocyte activation, the selected genes included: *C4b*, *C3*, *Serpina3n*, *Cxcl10*, *Gfap*, *Vim*, *Il18*, *Hif3*. Microglial activation genes included: *B2m*, *Trem2*, *Ccl2*, *Apoe*, *Axl*, *Itgax*, *Cd9*, *C1qa*, *C1qc*, *Lyz2*, and *Ctss*. The genes used for calculating the oligodendrocyte inflammation score were *C4b*, *I133*, and *Il18*. Genes utilized for estimating astrocyte A1 phenotype scores were: *C3*, *H2-T23*, *Serping1*, *H2-D1*, *Gbp2*, *Fkbp5*, *Psmb8*, *Srgn*, and *Amigo2*. The genes utilized for estimating astrocytes’ A2 phenotype scores were: *Clcf1*, *Tgm1*, *Ptx3*, *S100a10*, *Sphk1*, *Cd109*, *Ptgs2*, *Emp1*, *Slc10a6*, *Tm4sf1*, *B3gnt5*, *Cd14*, and *Stat3*.

Cell cycle phase was determined by score_genes_cell_cycle function in Scanpy. Gene references for cell cycle phase assignment are detailed in **Table S8**.

#### Gene module analysis within the lesion

To examine the gene modules associated with ICH pathology, we first used anatomical classification to identify the lesion area. We then performed a gene module analysis on this area. For the identification of functional gene modules, we utilized the Hotspot Python package^57^. The normalized gene expression matrix were used as the input. From this normalized data, we constructed a k-nearest neighbor graph of genes, utilizing the ’create_knn_graph’ function. The parameters set for this function were: ’n_neighbors = 10’. Following this, we retained the top 3000 genes that demonstrated significant autocorrelation, adhering to a False Discovery Rate (FDR) of less than 0.05. Identifying the modules use the ’create_modules’ function with the parameters: ’min_gene_threshold = 10’ and ’fdr_threshold = 0.05’. We used the enrichGO function from the clusterProfiler R package^109^ to annotate the identified gene modules. EnrichGO function was applied to each gene module list with the parameters: pvalueCutoff = 0.05, qvalueCutoff = 0.05, pAdjustMethod = "BH", ont = "ALL".

The gene module scores for other cells were calculated using scanpy.tl.score_genes function^106^. The score was the average expression of a set of genes subtracted with the average expression of a reference set of genes. The reference set was randomly sampled from the gene pool for each binned expression value.

#### Gene regulatory network construction

The Single-Cell rEgulatory Network Inference and Clustering (SCENIC) analysis was employed to uncover the gene regulatory network (GRN). This analysis was carried out using the latest version of pySCENIC (v0.12.1)^77^, which is a highly

efficient implementation of the SCENIC pipeline in Python. To define the search space around the transcription start site (TSS), gene-motif rankings were utilized based on sequences located 500 bp upstream or 100 bp downstream of the TSS. The motif database (mc9nr) was applied in the RcisTarget and GRNBoost algorithms to infer the gene regulatory networks.

#### Spatial gradient gene analysis

In this study, we conducted a spatial gradient gene analysis. Specifically, we first identified the lesion center and calculated the distance between each cell and the center. We then implemented generalized linear regression to establish a quantitative relationship between gene expression and distance. This analytic approach allowed us to efficiently and accurately decipher the spatial gene expression patterns within the lesion.

#### The identification of an injury-associated cell subclasses

To identify cell subclasses enriched in the injured hemisphere after ICH, we employed a systematic spatial analysis. Brain sections were digitally divided into a grid of 10x10 μm squares, allowing calculation of each hemisphere’s total area. Cell density (cells per unit area) for each subclass was then independently quantified in the left and right hemispheres. Subsequently, paired Wilcoxon tests were performed on each subclass to assess statistically significant differences (p-value < 0.05) in density between hemispheres. Cell subclasses exhibiting significant lateralization were identified as injury-associated.

### Immunofluorescence

After blocking with 3% bovine serum albumin, the primary antibodies were diluted in the antibody dilution buffer and incubated with the slices overnight at 4°C. The secondary antibodies were then diluted in the antibody dilution buffer and incubated with the slices for 30 min at room temperature. Finally, the sections were incubated with DAPI for 10 min to stain the nuclei. The primary antibodies used were CD68 (1:100, #ab283654, Abcam, Cambridge, MA, USA) and BODIPY (1mg/mL, #D3922, Thermo Fisher Scientific, Shanghai, China). Immunostained slices were imaged with captured by Leica Microsystems CMS GmbH Emst-Leitz-Str.17-37.

### Fluorescence in situ hybridization

The process of spatial Fluorescence *in situ* hybridization (FISH) involves designing highly efficient and specific target sequences using specific targeting probes provided by FISH Ltd. First, the mouse brain tissues are fixed, dehydrated, and paraffin-embedded and then sectioned into slices of 3 μm using a paraffin microtome. Afterward, the samples were treated with 4% paraformaldehyde and covered with a reaction chamber to perform the following reactions. The samples undergo dehydration and denaturation using 100% methanol. Subsequently, we added the hybridization buffer with specific targeting probes to the chamber, and the samples were incubated at 37□ overnight. The ligase reaction system was discarded, and the samples were thoroughly washed 3 times with phosphate-buffered saline with Tween 20 (PBST). The samples undergo rolling circle amplification by Phi29 DNA polymerase at 30□ overnight. Finally, fluorescent detection probes in the hybridization buffer were applied, and the samples were dehydrated with an ethanol series and mounted with a mounting medium. This process allows the damaged region of the sample to be imaged, providing valuable information on the spatial location of DNA. The FISH images were captured by Leica THUNDER Imaging Systems.

## Data and code availability

All data generated in this study are freely accessible in Spatial Transcript Omics DataBase (STOmics DB) under accession code sts0000122. All other data are in the main paper or the supplementary materials. Processed data can be interactively explored and downloaded from https://db.cngb.org/stomics/stmich. All the codes are publicly available and the source is annotated in the text and/or in the STAR Methods. Any additional information required to reanalyze the data reported in this paper is available from the lead contact upon request.

**Figure S1 Cell segmentation and quality control workflow, related to** Figure 1

(A) Cell segmentation process: In the upper part, each pixel represents a DNA nanoball (DNB) on the Stereo-seq chip, with 500 nanometers between each DNB. The color intensity indicates the number of RNA transcripts captured per DNB, with darker colors indicating higher transcript counts. The lower part shows cell nuclei stained from the same location, with each white point representing a nucleus. The rightmost image shows the merged DNBs and cell nuclei, illustrating the cell segmentation results. The yellow lines indicate the boundaries of each cell.

(B) The distribution of nuclear-localized RNA (*Malat1* and *Neat1*) and cytoplasmic RNA (mitochondria) in the sampled cells.

(C) Quality control and annotation workflow of Stereo-seq.

(D) Spatial visualization of primary cell types (left) along with corresponding markers (middle) and comparison with Allen Mouse Brain In Situ Hybridization (ISH) data (right).

**Figure S2 Marker gene expression and UMAP visualization of cell subclasses, related to** Figure 1

(A-N) Average marker gene expression after z-score transformation of cell subclasses (left), UMAP visualization of cell subclasses (middle), spatial visualization of cell subclasses (right).

**Figure S3 The activity of canonical signaling pathways**

(A) The activity of canonical signaling pathways at various time intervals.

**Figure S4 Anatomical partitioning, related to** Figure 2

(A) The alignment between molecularly defined tissue regions (left, colored based on molecular tissue region identities) and anatomically defined tissue regions (right, colored based on anatomically defined tissue regions). Each dot in the left hemisphere represents a cell. The anatomical definition of the right hemisphere comes from the Allen Brain Atlas. The abbreviation of the tissue region is consistent with the Allen Mouse Brain Reference Map. L2/3, cortical layer 2 and cortical layer 3; L4, cortical layer 4; L5, cortical layer 5; L6, cortical layer 6; lesion_1, lesion 1; lesion_2, lesion 2; Injured_STR_1, injured striatum 1; Injured_STR_2, injured striatum 2; Injured_STR_3, injured striatum 3; DG, dentate gyrus; FT, fiber tracts; HY, hypothalamus; PIR, piriform area; STRd, striatum dorsal region; STRv, striatum ventral region; TH, thalamus; VS, ventricular systems; isl,islm, islands of calleja and major island of calleja; PAL, pallidum. RT, reticular nucleus of the thalamus; LSX, lateral septal complex; sAMY, striatum-like amygdalar nuclei; CTXsp, cortical subplate;IG,Induseum griseum.

(B) Molecularly defined tissue regions across all slices, colored by defined regions.

**Figure S5 Spatial distribution and transcriptional signatures of glial activation, related to** Figure 3

(A) Spatial visualization of activation scores across different timepoints post-injury, for three types of glial cells: microglia (MGL), astrocytes (AC), and oligodendrocytes (OLG). The color intensity corresponds to the degree of activation, with red indicating the highest activation score and blue representing the lowest.

(B) Spatial visualization of genes specifically activated in three types of glial cells. Each plot illustrates the spatial distribution of select genes within their respective glial cell types at designated time points. Background cells are represented in yellow to highlight the gene expression in specific glial cell type. Specifically, *Mitf* is visualized in microglia, *Ptx3* and *Hbegf* in astrocytes, Gpd1 and *Cdkn1a* in oligodendrocytes.

(C) UpSet plot delineates the overlap of differentially expressed genes across three types of activated glial cells at various timepoints. The differentially expressed genes were identified by comparing each activated glial cell subclasses at each timepoint to its corresponding naive state. The horizontal bar chart on the bottom left displays the total number of differentially expressed genes within each dataset, while the vertical bar chart on the upper right illustrates the intersections between these datasets (with black dots indicating the presence of an intersection).

**Figure S6 Transcriptional signatures of glial activation, related to** Figure 3

(A) Spatial visualization of co-activated genes across three activated glial cells. Background cells are represented in yellow to highlight the gene expression in three glial cell type.

(B) Spatial visualization of genes that are upregulated specifically in astrocytes and oligodendrocytes. Changes in gene expression levels are shown for only these two cell types, with the other cell types showing the background in yellow.

**Figure S7 Functional diversity of astrocytes, related to** Figure 3

(A) Dot plot depicts the temporal expression patterns of genes *C3*, *Igfbp5*, and *Igfbp2* in activated astrocytes (AC_5) from the naive state through various post-ICH stages (Day 1 to Day 28). The color intensity represents the mean expression level within the group, while the dot size indicates the percentage of cells expressing the gene in the group.

(B) Spatial visualization of *C3*, *Igfbp5*, and *Igfbp2,* color by gene expression level.

**Figure S8 Additional gene modules, related to** Figure 4

(A) & (C) Spatial visualization of gene modules, color by module scores. (A) delineates gene module within normal function, and (C) focuses on gene modules with lesion. Key: ’M’ denotes modules.

(B) & (D) Gene ontology (go) enrichment analysis of gene modules from (A) & (C). X-axis: –log10 (p-adjust) score; Y-axis: GO terms. Only top 3 terms (p-adjust < 0.01) are listed for each gene module.

**Figure S9 Temporal characteristics of the lesion, related to** Figure 4

(A) Heatmap depicting the average expression of genes of gene module 42 across different time points within the lesion.

(B) Heatmap illustrating the scores of different cell subclasses from gene module 42 at different time points within the lesion.

(C) The distribution of cell cycle phases within all cells at various timepoints, from the naive state through subsequent days post-ICH (day 1 to day 28).

(D) Similar to (B), but for gene module 37.

(E) CV/LFB stained sections at different times.

(F) Spatial visualization of macrophage subclasses in the lesion.

**Figure S10 Visualization of genes related to the pathological process of intracerebral hemorrhage, related to** Figure 5

(A) The mean expression of scavenger receptor in different time points within the lesion.

(B) The left side of the figure presents spatial gene expression maps of phagocytosis-related genes (*Hmox1*, *Cd68*, *Ctsl*) within the lesion on day 7 post-ICH. The right side of the figure displays the mean expression of these phagocytosis-related genes in different cell types within the lesion on day 7 post-ICH. Additionally, genes with various functions are shown in (C) Iron metabolism, (D) Oxidation and antioxidation,

(E) Lipid metabolism and cholesterol transport, (F) Extracellular matrix, (G) Growth factor, (H) Neural repair.

**Figure S11 Molecular changes in different brain regions after intracerebral hemorrhage, related to** Figure 6

(A) The number of differentially expressed genes in various ipsilateral brain regions at different time points following ICH compared to the contralateral group.

Significant genes were identified based on specific criteria: logfoldchange≥1, scores > 5, and p-adjust<0.001.

(B) The heatmap illustrates the average expression levels of genes that are differentially expressed across cortical regions at various time points after ICH, in comparison to naïve controls. Columns represent distinct cortical areas at specific time points post-ICH, while rows correspond to individual genes.

(C) Bubble plots depicts the GO enrichment analysis results for genes differentially expressed in the L4 cortical layer at successive time points following ICH.

(D) Spatial visualization of *Dlx1, Dlx2, Sox4, and Sox11*.

(E) Spatial visualization of *Dcx*.

(F) Spatial visualization of *Serpina3n, C4b, Gfap, and Klk6*.

## Supplementary information

Table S1: This table includes quality control information for each slice used in the study.

Table S2: Classic gene markers used for cell annotation are listed in this table.

Table S3: This table lists differentially expressed genes for each cell subclass.

Table S4: Differentially expressed genes in the three glia cell types at different time points, the differentially expressed genes were identified by comparing each activated glial cell subclass at each timepoint to its corresponding naive state.

Table S5: This table lists gene modules, along with the GO enrichment analysis results for these gene modules.

Table S6: This table lists the differentially expressed genes identified through two comparison methods, including comparisons between the post-ICH groups and the naive group, as well as comparisons between the ipsilateral and contralateral group.

Table S7: The results of GO enrichment analysis for the differentially expressed genes of different region are presented in this table. It includes comparisons between the post-ICH group and the naive group, as well as comparisons between the ipsilateral and contralateral groups.

Table S8: This table lists cell cycle phase genes

## Reference

1. Xue, M., and Yong, V.W. (2020). Neuroinflammation in intracerebral haemorrhage: immunotherapies with potential for translation. The Lancet Neurology 19, 1023–1032. 10.1016/S1474-4422(20)30364-1.

2. Ren, H., Han, R., Chen, X., Liu, X., Wan, J., Wang, L., Yang, X., and Wang, J. (2020). Potential therapeutic targets for intracerebral hemorrhage-associated inflammation: An update. J Cereb Blood Flow Metab 40, 1752–1768. 10.1177/0271678X20923551.

3. Hanley, D.F., Thompson, R.E., Rosenblum, M., Yenokyan, G., Lane, K., McBee, N., Mayo, S.W., Bistran-Hall, A.J., Gandhi, D., Mould, W.A., et al. (2019). Efficacy and safety of minimally invasive surgery with thrombolysis in intracerebral haemorrhage evacuation (MISTIE III): a randomised, controlled, open-label, blinded endpoint phase 3 trial. Lancet 393, 1021–1032. 10.1016/S0140-6736(19)30195-3.

4. Zhang, Z., Xu, W., Sheng, H., Huang, L., Zhang, J., Zhang, L., Wang, L., Wang, J., Ren, X., Jiang, C., et al. (2023). Hematoma clearance by reactive microglia after intracerebral hemorrhage. Gene & Protein in Disease 2, 336. 10.36922/gpd.336.

5. Lan, X., Han, X., Liu, X., and Wang, J. (2019). Inflammatory responses after intracerebral hemorrhage: From cellular function to therapeutic targets. J Cereb Blood Flow Metab 39, 184–186. 10.1177/0271678X18805675.

6. Liddelow, S.A., Guttenplan, K.A., Clarke, L.E., Bennett, F.C., Bohlen, C.J., Schirmer, L., Bennett, M.L., Münch, A.E., Chung, W.-S., Peterson, T.C., et al. (2017). Neurotoxic reactive astrocytes are induced by activated microglia. Nature 541, 481–487. 10.1038/nature21029.

7. Greenhalgh, A.D., David, S., and Bennett, F.C. (2020). Immune cell regulation of glia during CNS injury and disease. Nat Rev Neurosci 21, 139–152. 10.1038/s41583-020-0263-9.

8. Lan, X., Han, X., Li, Q., Yang, Q.-W., and Wang, J. (2017). Modulators of microglial activation and polarization after intracerebral haemorrhage. Nat Rev Neurol 13, 420–433. 10.1038/nrneurol.2017.69.

9. Pekny, M., Wilhelmsson, U., Tatlisumak, T., and Pekna, M. (2019). Astrocyte activation and reactive gliosis-A new target in stroke? Neurosci Lett 689, 45–55. 10.1016/j.neulet.2018.07.021.

10. Zhu, H., Wang, Z., Yu, J., Yang, X., He, F., Liu, Z., Che, F., Chen, X., Ren, H., Hong, M., et al. (2019). Role and mechanisms of cytokines in the secondary brain injury after intracerebral hemorrhage. Prog Neurobiol 178, 101610. 10.1016/j.pneurobio.2019.03.003.

11. Song, D., Yeh, C.-T., Wang, J., and Guo, F. (2022). Perspectives on the mechanism of pyroptosis after intracerebral hemorrhage. Front Immunol 13, 989503. 10.3389/fimmu.2022.989503.

12. Li, Q., Lan, X., Han, X., Durham, F., Wan, J., Weiland, A., Koehler, R.C., and Wang, J. (2021). Microglia-derived interleukin-10 accelerates post-intracerebral hemorrhage hematoma clearance by regulating CD36. Brain Behav Immun 94, 437–457. 10.1016/j.bbi.2021.02.001.

13. 13. Chang, C.-F., Goods, B.A., Askenase, M.H., Hammond, M.D., Renfroe, S.C., Steinschneider, A.F., Landreneau, M.J., Ai, Y., Beatty, H.E., Da Costa, L.H.A., et al. (2017). Erythrocyte efferocytosis modulates macrophages towards recovery after intracerebral hemorrhage. Journal of Clinical Investigation 128, 607–624. 10.1172/JCI95612.

14. Shi, E., Shi, K., Qiu, S., Sheth, K.N., Lawton, M.T., and Ducruet, A.F. (2019). Chronic inflammation, cognitive impairment, and distal brain region alteration following intracerebral hemorrhage. FASEB j. 33, 9616–9626. 10.1096/fj.201900257R.

15. Liang, C., Liu, L., Bao, S., Yao, Z., Bai, Q., Fu, P., Liu, X., Zhang, J.H., and Wang, G. (2023). Neuroprotection by Nrf2 via modulating microglial phenotype and phagocytosis after intracerebral hemorrhage. Heliyon 9, e13777. 10.1016/j.heliyon.2023.e13777.

16. Li, H., Ghorbani, S., Zhang, R., Ebacher, V., Stephenson, E.L., Keough, M.B., Yong, V.W., and Xue, M. (2023). Prominent elevation of extracellular matrix molecules in intracerebral hemorrhage. Front Mol Neurosci 16, 1251432. 10.3389/fnmol.2023.1251432.

17. Liu, Y., Qi, L., Li, Z., Yong, V.W., and Xue, M. (2023). Crosstalk Between Matrix Metalloproteinases and Their Inducer EMMPRIN/CD147: a Promising Therapeutic Target for Intracerebral Hemorrhage. Transl Stroke Res. 10.1007/s12975-023-01225-6.

18. Puy, L., Parry-Jones, A.R., Sandset, E.C., Dowlatshahi, D., Ziai, W., and Cordonnier, C. (2023). Intracerebral haemorrhage. Nat Rev Dis Primers 9, 1–18. 10.1038/s41572-023-00424-7.

19. Ironside, N., Chen, C.-J., Ding, D., Mayer, S.A., and Connolly, E.S. (2019). Perihematomal Edema After Spontaneous Intracerebral Hemorrhage. Stroke 50, 1626–1633. 10.1161/STROKEAHA.119.024965.

20. Chen, A. Spatiotemporal transcriptomic atlas of mouse organogenesis using DNA nanoball-patterned arrays. OPEN ACCESS, 38.

21. Carey, J.R., Evans, C.D., Anderson, D.C., Bhatt, E., Nagpal, A., Kimberley, T.J., and Pascual-Leone, A. (2008). Safety of 6-Hz primed low-frequency rTMS in stroke. Neurorehabil Neural Repair 22, 185–192. 10.1177/1545968307305458.

22. Li, Q., Han, X., Lan, X., Gao, Y., Wan, J., Durham, F., Cheng, T., Yang, J., Wang, Z., Jiang, C., et al. (2017). Inhibition of neuronal ferroptosis protects hemorrhagic brain. JCI Insight 2, e90777. 10.1172/jci.insight.90777.

23. Jia, P., He, J., Li, Z., Wang, J., Jia, L., Hao, R., Lai, J., Zang, W., Chen, X., and Wang, J. (2021). Profiling of Blood-Brain Barrier Disruption in Mouse Intracerebral Hemorrhage Models: Collagenase Injection vs. Autologous Arterial Whole Blood Infusion. Front Cell Neurosci 15, 699736. 10.3389/fncel.2021.699736.

24. Wang, J. (2010). Preclinical and clinical research on inflammation after intracerebral hemorrhage. Prog Neurobiol 92, 463–477. 10.1016/j.pneurobio.2010.08.001.

25. Stringer, C., Wang, T., Michaelos, M., and Pachitariu, M. (2021). Cellpose: a generalist algorithm for cellular segmentation. Nat Methods 18, 100–106. 10.1038/s41592-020-01018-x.

26. Lopez, R., Regier, J., Cole, M.B., Jordan, M.I., and Yosef, N. (2018). Deep generative modeling for single-cell transcriptomics. Nat Methods 15, 1053–1058. 10.1038/s41592-018-0229-2.

27. Xu, C., Lopez, R., Mehlman, E., Regier, J., Jordan, M.I., and Yosef, N. (2021). Probabilistic harmonization and annotation of single-cell transcriptomics data with deep generative models. Mol Syst Biol 17, e9620. 10.15252/msb.20209620.

28. Shi, H., He, Y., Zhou, Y., Huang, J., Maher, K., Wang, B., Tang, Z., Luo, S., Tan, P., Wu, M., et al. (2023). Spatial atlas of the mouse central nervous system at molecular resolution. Nature 622, 552–561. 10.1038/s41586-023-06569-5.

29. Schubert, M., Klinger, B., Klünemann, M., Sieber, A., Uhlitz, F., Sauer, S., Garnett, M.J., Blüthgen, N., and Saez-Rodriguez, J. (2018). Perturbation-response genes reveal signaling footprints in cancer gene expression. Nat Commun 9, 20. 10.1038/s41467-017-02391-6.

30. Holland, C.H., Szalai, B., and Saez-Rodriguez, J. (2020). Transfer of regulatory knowledge from human to mouse for functional genomics analysis. Biochim Biophys Acta Gene Regul Mech 1863, 194431. 10.1016/j.bbagrm.2019.194431.

31. 31. Ben Haim, L., Ceyzériat, K., Carrillo-de Sauvage, M.A., Aubry, F., Auregan, G., Guillermier, M., Ruiz, M., Petit, F., Houitte, D., Faivre, E., et al. (2015). The JAK/STAT3 pathway is a common inducer of astrocyte reactivity in Alzheimer’s and Huntington’s diseases. J Neurosci 35, 2817–2829. 10.1523/JNEUROSCI.3516-14.2015.

32. Zheng, J., Wu, H., Wang, X., Zhang, G., Lu, J., Xu, W., Xu, S., Fang, Y., Zhang, A., Shao, A., et al. (2023). Temporal dynamics of microglia-astrocyte interaction in neuroprotective glial scar formation after intracerebral hemorrhage. Journal of Pharmaceutical Analysis. 10.1016/j.jpha.2023.02.007.

33. He, Y., Tang, X., Huang, J., Ren, J., Zhou, H., Chen, K., Liu, A., Shi, H., Lin, Z., Li, Q., et al. (2021). ClusterMap for multi-scale clustering analysis of spatial gene expression. Nat Commun 12, 5909. 10.1038/s41467-021-26044-x.

34. Kang, M., and Yao, Y. (2019). Oligodendrocytes in intracerebral hemorrhage. CNS Neurosci Ther 25, 1075–1084. 10.1111/cns.13193.

35. Clarke, L.E., Liddelow, S.A., Chakraborty, C., Münch, A.E., Heiman, M., and Barres, B.A. (2018). Normal aging induces A1-like astrocyte reactivity. Proceedings of the National Academy of Sciences 115, E1896–E1905. 10.1073/pnas.1800165115.

36. Hammond, T.R., Dufort, C., Dissing-Olesen, L., Giera, S., Young, A., Wysoker, A., Walker, A.J., Gergits, F., Segel, M., Nemesh, J., et al. (2019). Single-Cell RNA Sequencing of Microglia throughout the Mouse Lifespan and in the Injured Brain Reveals Complex Cell-State Changes. Immunity 50, 253–271.e6. 10.1016/j.immuni.2018.11.004.

37. Keren-Shaul, H., Spinrad, A., Weiner, A., Matcovitch-Natan, O., Dvir-Szternfeld, R., Ulland, T.K., David, E., Baruch, K., Lara-Astaiso, D., Toth, B., et al. (2017). A Unique Microglia Type Associated with Restricting Development of Alzheimer’s Disease. Cell 169, 1276–1290.e17. 10.1016/j.cell.2017.05.018.

38. Dolan, M.-J., Therrien, M., Jereb, S., Kamath, T., Gazestani, V., Atkeson, T., Marsh, S.E., Goeva, A., Lojek, N.M., Murphy, S., et al. (2023). Exposure of iPSC-derived human microglia to brain substrates enables the generation and manipulation of diverse transcriptional states in vitro. Nat Immunol 24, 1382–1390. 10.1038/s41590-023-01558-2.

39. Shindo, A., Maki, T., Mandeville, E.T., Liang, A.C., Egawa, N., Itoh, K., Itoh, N., Borlongan, M., Holder, J.C., Chuang, T.T., et al. (2016). Astrocyte-Derived Pentraxin 3 Supports Blood-Brain Barrier Integrity Under Acute Phase of Stroke. Stroke 47, 1094–1100. 10.1161/STROKEAHA.115.012133.

40. 40. Puschmann, T.B., Zandén, C., Lebkuechner, I., Philippot, C., de Pablo, Y., Liu, J., and Pekny, M. (2014). HB-EGF affects astrocyte morphology, proliferation, differentiation, and the expression of intermediate filament proteins. J Neurochem 128, 878–889. 10.1111/jnc.12519.

41. Ni, W., Zheng, M., Xi, G., Keep, R.F., and Hua, Y. (2015). Role of lipocalin-2 in brain injury after intracerebral hemorrhage. J Cereb Blood Flow Metab 35, 1454– 1461. 10.1038/jcbfm.2015.52.

42. 42. Tomás-Camardiel, M., Venero, J.L., de Pablos, R.M., Rite, I., Machado, A., and Cano, J. (2004). In vivo expression of aquaporin-4 by reactive microglia. J Neurochem 91, 891–899. 10.1111/j.1471-4159.2004.02759.x.

43. Liu, C., Zhao, X.-M., Wang, Q., Du, T.-T., Zhang, M.-X., Wang, H.-Z., Li, R.-P., Liang, K., Gao, Y., Zhou, S.-Y., et al. (2023). Astrocyte-derived SerpinA3N promotes neuroinflammation and epileptic seizures by activating the NF-κB signaling pathway in mice with temporal lobe epilepsy. J Neuroinflammation 20, 161. 10.1186/s12974-023-02840-8.

44. Hou, J., Zhou, Y., Cai, Z., Terekhova, M., Swain, A., Andhey, P.S., Guimaraes, R.M., Ulezko Antonova, A., Qiu, T., Sviben, S., et al. (2023). Transcriptomic atlas and interaction networks of brain cells in mouse CNS demyelination and remyelination. Cell Rep 42, 112293. 10.1016/j.celrep.2023.112293.

45. Pekny, M., and Pekna, M. (2014). Astrocyte reactivity and reactive astrogliosis: costs and benefits. Physiol Rev 94, 1077–1098. 10.1152/physrev.00041.2013.

46. Baer, A.S., Syed, Y.A., Kang, S.U., Mitteregger, D., Vig, R., Ffrench-Constant, C., Franklin, R.J.M., Altmann, F., Lubec, G., and Kotter, M.R. (2009). Myelin-mediated inhibition of oligodendrocyte precursor differentiation can be overcome by pharmacological modulation of Fyn-RhoA and protein kinase C signalling. Brain 132, 465–481. 10.1093/brain/awn334.

47. Zhang, Z.-H., Ma, F.-F., Zhang, H., and Xu, X.-H. (2017). MARCKS is Necessary for Oligodendrocyte Precursor Cell Maturation. Neurochem Res 42, 2933– 2939. 10.1007/s11064-017-2324-7.

48. 48. Sirko, S., Irmler, M., Gascón, S., Bek, S., Schneider, S., Dimou, L., Obermann, J., De Souza Paiva, D., Poirier, F., Beckers, J., et al. (2015). Astrocyte reactivity after brain injury-: The role of galectins 1 and 3. Glia 63, 2340–2361. 10.1002/glia.22898.

49. Walter, H.J., Berry, M., Hill, D.J., and Logan, A. (1997). Spatial and temporal changes in the insulin-like growth factor (IGF) axis indicate autocrine/paracrine actions of IGF-I within wounds of the rat brain. Endocrinology 138, 3024–3034. 10.1210/endo.138.7.5284.

50. Zamanian, J.L., Xu, L., Foo, L.C., Nouri, N., Zhou, L., Giffard, R.G., and Barres, B.A. (2012). Genomic Analysis of Reactive Astrogliosis. J. Neurosci. 32, 6391–6410. 10.1523/JNEUROSCI.6221-11.2012.

51. Escartin, C., Galea, E., Lakatos, A., O’Callaghan, J.P., Petzold, G.C., Serrano-Pozo, A., Steinhäuser, C., Volterra, A., Carmignoto, G., Agarwal, A., et al. (2021). Reactive astrocyte nomenclature, definitions, and future directions. Nat Neurosci 24, 312–325. 10.1038/s41593-020-00783-4.

52. Miller, S.J. (2018). Astrocyte Heterogeneity in the Adult Central Nervous System. Front Cell Neurosci 12, 401. 10.3389/fncel.2018.00401.

53. Hartmann, K., Sepulveda-Falla, D., Rose, I.V.L., Madore, C., Muth, C., Matschke, J., Butovsky, O., Liddelow, S., Glatzel, M., and Krasemann, S. (2019). Complement 3+-astrocytes are highly abundant in prion diseases, but their abolishment led to an accelerated disease course and early dysregulation of microglia. Acta Neuropathol Commun 7, 83. 10.1186/s40478-019-0735-1.

54. Nold, V., Richter, N., Hengerer, B., Kolassa, I.-T., and Allers, K.A. (2021). FKBP5 polymorphisms induce differential glucocorticoid responsiveness in primary CNS cells - First insights from novel humanized mice. Eur J Neurosci 53, 402–415. 10.1111/ejn.14999.

55. Ceyzériat, K., Ben Haim, L., Denizot, A., Pommier, D., Matos, M., Guillemaud, O., Palomares, M.-A., Abjean, L., Petit, F., Gipchtein, P., et al. (2018). Modulation of astrocyte reactivity improves functional deficits in mouse models of Alzheimer’s disease. Acta Neuropathol Commun 6, 104. 10.1186/s40478-018-0606-1.

56. Klemens, J., Ciurkiewicz, M., Chludzinski, E., Iseringhausen, M., Klotz, D., Pfankuche, V.M., Ulrich, R., Herder, V., Puff, C., Baumgärtner, W., et al. (2019). Neurotoxic potential of reactive astrocytes in canine distemper demyelinating leukoencephalitis. Sci Rep 9, 11689. 10.1038/s41598-019-48146-9.

57. DeTomaso, D., and Yosef, N. (2021). Hotspot identifies informative gene modules across modalities of single-cell genomics. Cell Systems 12, 446–456.e9. 10.1016/j.cels.2021.04.005.

58. Park, D.S., Levine, B., Ferrari, G., and Greene, L.A. (1997). Cyclin dependent kinase inhibitors and dominant negative cyclin dependent kinase 4 and 6 promote survival of NGF-deprived sympathetic neurons. J Neurosci 17, 8975–8983. 10.1523/JNEUROSCI.17-23-08975.1997.

59. Park, D.S., Morris, E.J., Stefanis, L., Troy, C.M., Shelanski, M.L., Geller, H.M., and Greene, L.A. (1998). Multiple pathways of neuronal death induced by DNA-damaging agents, NGF deprivation, and oxidative stress. J Neurosci 18, 830– 840. 10.1523/JNEUROSCI.18-03-00830.1998.

60. Giovanni, A., Wirtz-Brugger, F., Keramaris, E., Slack, R., and Park, D.S. (1999). Involvement of cell cycle elements, cyclin-dependent kinases, pRb, and E2F x DP, in B-amyloid-induced neuronal death. J Biol Chem 274, 19011–19016. 10.1074/jbc.274.27.19011.

61. Herrup, K., and Yang, Y. (2007). Cell cycle regulation in the postmitotic neuron: oxymoron or new biology? Nat Rev Neurosci 8, 368–378. 10.1038/nrn2124.

62. Stolt, C.C., Rehberg, S., Ader, M., Lommes, P., Riethmacher, D., Schachner, M., Bartsch, U., and Wegner, M. (2002). Terminal differentiation of myelin-forming oligodendrocytes depends on the transcription factor Sox10. Genes Dev 16, 165–170. 10.1101/gad.215802.

63. Hartwig, C., Veske, A., Krejcova, S., Rosenberger, G., and Finckh, U. (2005). Plexin B3 promotes neurite outgrowth, interacts homophilically, and interacts with Rin. BMC Neurosci 6, 53. 10.1186/1471-2202-6-53.

64. Wan, J., Zhang, G., Li, X., Qiu, X., Ouyang, J., Dai, J., and Min, S. (2021). Matrix Metalloproteinase 3: A Promoting and Destabilizing Factor in the Pathogenesis of Disease and Cell Differentiation. Front Physiol 12, 663978. 10.3389/fphys.2021.663978.

65. Hanayama, R., Tanaka, M., Miwa, K., Shinohara, A., Iwamatsu, A., and Nagata, S. (2002). Identification of a factor that links apoptotic cells to phagocytes. Nature 417, 182–187. 10.1038/417182a.

66. Hanayama, R., Tanaka, M., Miyasaka, K., Aozasa, K., Koike, M., Uchiyama, Y., and Nagata, S. (2004). Autoimmune disease and impaired uptake of apoptotic cells in MFG-E8-deficient mice. Science 304, 1147–1150. 10.1126/science.1094359.

67. Laplante, P., Brillant-Marquis, F., Brissette, M.-J., Joannette-Pilon, B., Cayrol, R., Kokta, V., and Cailhier, J.-F. (2017). MFG-E8 Reprogramming of Macrophages Promotes Wound Healing by Increased bFGF Production and Fibroblast Functions. J Invest Dermatol 137, 2005–2013. 10.1016/j.jid.2017.04.030.

68. Wang, J., and Doré, S. (2007). Heme oxygenase-1 exacerbates early brain injury after intracerebral haemorrhage. Brain 130, 1643–1652. 10.1093/brain/awm095.

69. Qu, J., Chen, W., Hu, R., and Feng, H. (2016). The Injury and Therapy of Reactive Oxygen Species in Intracerebral Hemorrhage Looking at Mitochondria. Oxidative Medicine and Cellular Longevity 2016, e2592935. 10.1155/2016/2592935.

70. Duan, X., Wen, Z., Shen, H., Shen, M., and Chen, G. (2016). Intracerebral Hemorrhage, Oxidative Stress, and Antioxidant Therapy. Oxidative Medicine and Cellular Longevity 2016, e1203285. 10.1155/2016/1203285.

71. Robinson, S.R., Dang, T.N., Dringen, R., and Bishop, G.M. (2009). Hemin toxicity: a preventable source of brain damage following hemorrhagic stroke. Redox Report 14, 228–235. 10.1179/135100009X12525712409931.

72. Sherman, D.L., Tait, S., Melrose, S., Johnson, R., Zonta, B., Court, F.A., Macklin, W.B., Meek, S., Smith, A.J.H., Cottrell, D.F., et al. (2005). Neurofascins are required to establish axonal domains for saltatory conduction. Neuron 48, 737–742. 10.1016/j.neuron.2005.10.019.

73. Zhang, A., Desmazieres, A., Zonta, B., Melrose, S., Campbell, G., Mahad, D., Li, Q., Sherman, D.L., Reynolds, R., and Brophy, P.J. (2015). Neurofascin 140 is an embryonic neuronal neurofascin isoform that promotes the assembly of the node of Ranvier. J Neurosci 35, 2246–2254. 10.1523/JNEUROSCI.3552-14.2015.

74. Zonta, B., Desmazieres, A., Rinaldi, A., Tait, S., Sherman, D.L., Nolan, M.F., and Brophy, P.J. (2011). A critical role for Neurofascin in regulating action potential initiation through maintenance of the axon initial segment. Neuron 69, 945–956. 10.1016/j.neuron.2011.02.021.

75. Ma, X.-R., Zhu, X., Xiao, Y., Gu, H.-M., Zheng, S.-S., Li, L., Wang, F., Dong, Z.-J., Wang, D.-X., Wu, Y., et al. (2022). Restoring nuclear entry of Sirtuin 2 in oligodendrocyte progenitor cells promotes remyelination during ageing. Nat Commun 13, 1225. 10.1038/s41467-022-28844-1.

76. Lee, J., Gravel, M., Zhang, R., Thibault, P., and Braun, P.E. (2005). Process outgrowth in oligodendrocytes is mediated by CNP, a novel microtubule assembly myelin protein. J Cell Biol 170, 661–673. 10.1083/jcb.200411047.

77. Aibar, S., González-Blas, C.B., Moerman, T., Huynh-Thu, V.A., Imrichova, H., Hulselmans, G., Rambow, F., Marine, J.-C., Geurts, P., Aerts, J., et al. (2017). SCENIC: single-cell regulatory network inference and clustering. Nat Methods 14, 1083–1086. 10.1038/nmeth.4463.

78. Gao, M., Sossa, K., Song, L., Errington, L., Cummings, L., Hwang, H., Kuhl, D., Worley, P., and Lee, H.-K. (2010). A specific requirement of Arc/Arg3.1 for visual experience-induced homeostatic synaptic plasticity in mouse primary visual cortex. J Neurosci 30, 7168–7178. 10.1523/JNEUROSCI.1067-10.2010.

79. Das, S., Lituma, P.J., Castillo, P.E., and Singer, R.H. (2023). Maintenance of a short-lived protein required for long-term memory involves cycles of transcription and local translation. Neuron 111, 2051–2064.e6. 10.1016/j.neuron.2023.04.005.

80. Li, L., Yun, S.H., Keblesh, J., Trommer, B.L., Xiong, H., Radulovic, J., and Tourtellotte, W.G. (2007). Egr3, a synaptic activity regulated transcription factor that is essential for learning and memory. Mol Cell Neurosci 35, 76–88. 10.1016/j.mcn.2007.02.004.

81. Choy, F.C., Klarić, T.S., Leong, W.K., Koblar, S.A., and Lewis, M.D. (2016). Reduction of the neuroprotective transcription factor Npas4 results in increased neuronal necrosis, inflammation and brain lesion size following ischaemia. J Cereb Blood Flow Metab 36, 1449–1463. 10.1177/0271678X15606146.

82. Lin, Y., Bloodgood, B.L., Hauser, J.L., Lapan, A.D., Koon, A.C., Kim, T.-K., Hu, L.S., Malik, A.N., and Greenberg, M.E. (2008). Activity-dependent regulation of inhibitory synapse development by Npas4. Nature 455, 1198–1204. 10.1038/nature07319.

83. 83. Jeanneteau, F., Barrère, C., Vos, M., De Vries, C.J.M., Rouillard, C., Levesque, D., Dromard, Y., Moisan, M.-P., Duric, V., Franklin, T.C., et al. (2018). The Stress-Induced Transcription Factor NR4A1 Adjusts Mitochondrial Function and Synapse Number in Prefrontal Cortex. J Neurosci 38, 1335–1350. 10.1523/JNEUROSCI.2793-17.2017.

84. Codega, P., Silva-Vargas, V., Paul, A., Maldonado-Soto, A.R., Deleo, A.M., Pastrana, E., and Doetsch, F. (2014). Prospective identification and purification of quiescent adult neural stem cells from their in vivo niche. Neuron 82, 545–559. 10.1016/j.neuron.2014.02.039.

85. Hsieh, J. (2012). Orchestrating transcriptional control of adult neurogenesis. Genes Dev. 26, 1010–1021. 10.1101/gad.187336.112.

86. Cameron, E.G., Nahmou, M., Toth, A.B., Heo, L., Tanasa, B., Dalal, R., Yan, W., Nallagatla, P., Xia, X., Hay, S., et al. (2024). A molecular switch for neuroprotective astrocyte reactivity. Nature 626, 574–582. 10.1038/s41586-023-06935-3.

87. Zbesko, J.C., Stokes, J., Becktel, D.A., and Doyle, K.P. (2023). Targeting foam cell formation to improve recovery from ischemic stroke. Neurobiology of Disease 181, 106130. 10.1016/j.nbd.2023.106130.

88. Zhang, R., Dong, Y., Liu, Y., Moezzi, D., Ghorbani, S., Mirzaei, R., Lozinski, B.M., Dunn, J.F., Yong, V.W., and Xue, M. (2023). Enhanced liver X receptor signalling reduces brain injury and promotes tissue regeneration following experimental intracerebral haemorrhage: roles of microglia/macrophages. Stroke Vasc Neurol 8, 486–502. 10.1136/svn-2023-002331.

89. Sage, A.P., Tsiantoulas, D., Binder, C.J., and Mallat, Z. (2019). The role of B cells in atherosclerosis. Nat Rev Cardiol 16, 180–196. 10.1038/s41569-018-0106-9.

90. Tsiantoulas, D., Diehl, C.J., Witztum, J.L., and Binder, C.J. (2014). B cells and humoral immunity in atherosclerosis. Circ Res 114, 1743–1756. 10.1161/CIRCRESAHA.113.301145.

91. Arvidsson, A., Collin, T., Kirik, D., Kokaia, Z., and Lindvall, O. (2002). Neuronal replacement from endogenous precursors in the adult brain after stroke. Nat Med 8, 963–970. 10.1038/nm747.

92. Parent, J.M., Vexler, Z.S., Gong, C., Derugin, N., and Ferriero, D.M. (2002). Rat forebrain neurogenesis and striatal neuron replacement after focal stroke. Ann Neurol 52, 802–813. 10.1002/ana.10393.

93. Lagace, D.C. (2012). Does the endogenous neurogenic response alter behavioral recovery following stroke? Behavioural Brain Research 227, 426–432. 10.1016/j.bbr.2011.08.045.

94. Ohab, J.J., Fleming, S., Blesch, A., and Carmichael, S.T. (2006). A neurovascular niche for neurogenesis after stroke. J Neurosci 26, 13007–13016. 10.1523/JNEUROSCI.4323-06.2006.

95. Guzowski, J.F., Timlin, J.A., Roysam, B., McNaughton, B.L., Worley, P.F., and Barnes, C.A. (2005). Mapping behaviorally relevant neural circuits with immediate-early gene expression. Current Opinion in Neurobiology 15, 599–606. 10.1016/j.conb.2005.08.018.

96. Kim, S., Kim, H., and Um, J.W. (2018). Synapse development organized by neuronal activity-regulated immediate-early genes. Exp Mol Med 50, 1–7. 10.1038/s12276-018-0025-1.

97. Shepherd, J.D., and Bear, M.F. (2011). New views of Arc, a master regulator of synaptic plasticity. Nat Neurosci 14, 279–284. 10.1038/nn.2708.

98. Goto, A. (2022). Synaptic plasticity during systems memory consolidation. Neuroscience Research 183, 1–6. 10.1016/j.neures.2022.05.008.

99. Minatohara, K., Akiyoshi, M., and Okuno, H. (2015). Role of Immediate-Early Genes in Synaptic Plasticity and Neuronal Ensembles Underlying the Memory Trace. Front Mol Neurosci 8, 78. 10.3389/fnmol.2015.00078.

100. Byrns, C.N., Saikumar, J., and Bonini, N.M. (2021). Glial AP1 is activated with aging and accelerated by traumatic brain injury. Nat Aging 1, 585–597. 10.1038/s43587-021-00072-0.

101. Jia, P., Wang, J., Ren, X., He, J., Wang, S., Xing, Y., Chen, D., Zhang, X., Zhou, S., Liu, X., et al. (2023). An enriched environment improves long-term functional outcomes in mice after intracerebral hemorrhage by mechanisms that involve the Nrf2/BDNF/glutaminase pathway. J Cereb Blood Flow Metab 43, 694– 711. 10.1177/0271678X221135419.

102. Li, Q., Wan, J., Lan, X., Han, X., Wang, Z., and Wang, J. (2017). Neuroprotection of brain-permeable iron chelator VK-28 against intracerebral hemorrhage in mice. J Cereb Blood Flow Metab 37, 3110–3123. 10.1177/0271678X17709186.

103. Dobin, A., Davis, C.A., Schlesinger, F., Drenkow, J., Zaleski, C., Jha, S., Batut, P., Chaisson, M., and Gingeras, T.R. (2013). STAR: ultrafast universal RNA-seq aligner. Bioinformatics 29, 15–21. 10.1093/bioinformatics/bts635.

104. Pachitariu, M., and Stringer, C. (2022). Cellpose 2.0: how to train your own model. Nat Methods 19, 1634–1641. 10.1038/s41592-022-01663-4.

105. 105. Shi, H., He, Y., Zhou, Y., Huang, J., Wang, B., Tang, Z., Tan, P., Wu, M., Lin, Z., Ren, J., et al. (2022). Spatial Atlas of the Mouse Central Nervous System at Molecular Resolution (Neuroscience) 10.1101/2022.06.20.496914.

106. Wolf, F.A., Angerer, P., and Theis, F.J. (2018). SCANPY: large-scale single-cell gene expression data analysis. Genome Biology 19, 15. 10.1186/s13059-017-1382-0.

107. 107. Traag, V.A., Waltman, L., and van Eck, N.J. (2019). From Louvain to Leiden: guaranteeing well-connected communities. Sci Rep 9, 5233. 10.1038/s41598-019-41695-z.

108. Badia-i-Mompel, P., Vélez Santiago, J., Braunger, J., Geiss, C., Dimitrov, D., Müller-Dott, S., Taus, P., Dugourd, A., Holland, C.H., Ramirez Flores, R.O., et al. (2022). decoupleR: ensemble of computational methods to infer biological activities from omics data. Bioinformatics Advances 2, vbac016. 10.1093/bioadv/vbac016.

109. Wu, T., Hu, E., Xu, S., Chen, M., Guo, P., Dai, Z., Feng, T., Zhou, L., Tang, W., Zhan, L., et al. (2021). clusterProfiler 4.0: A universal enrichment tool for interpreting omics data. The Innovation 2, 100141. 10.1016/j.xinn.2021.100141.

