## Supplementary material for "Spatiotemporal transcriptomic maps of mouse intracerebral hemorrhage at single-cell resolution": Figure S

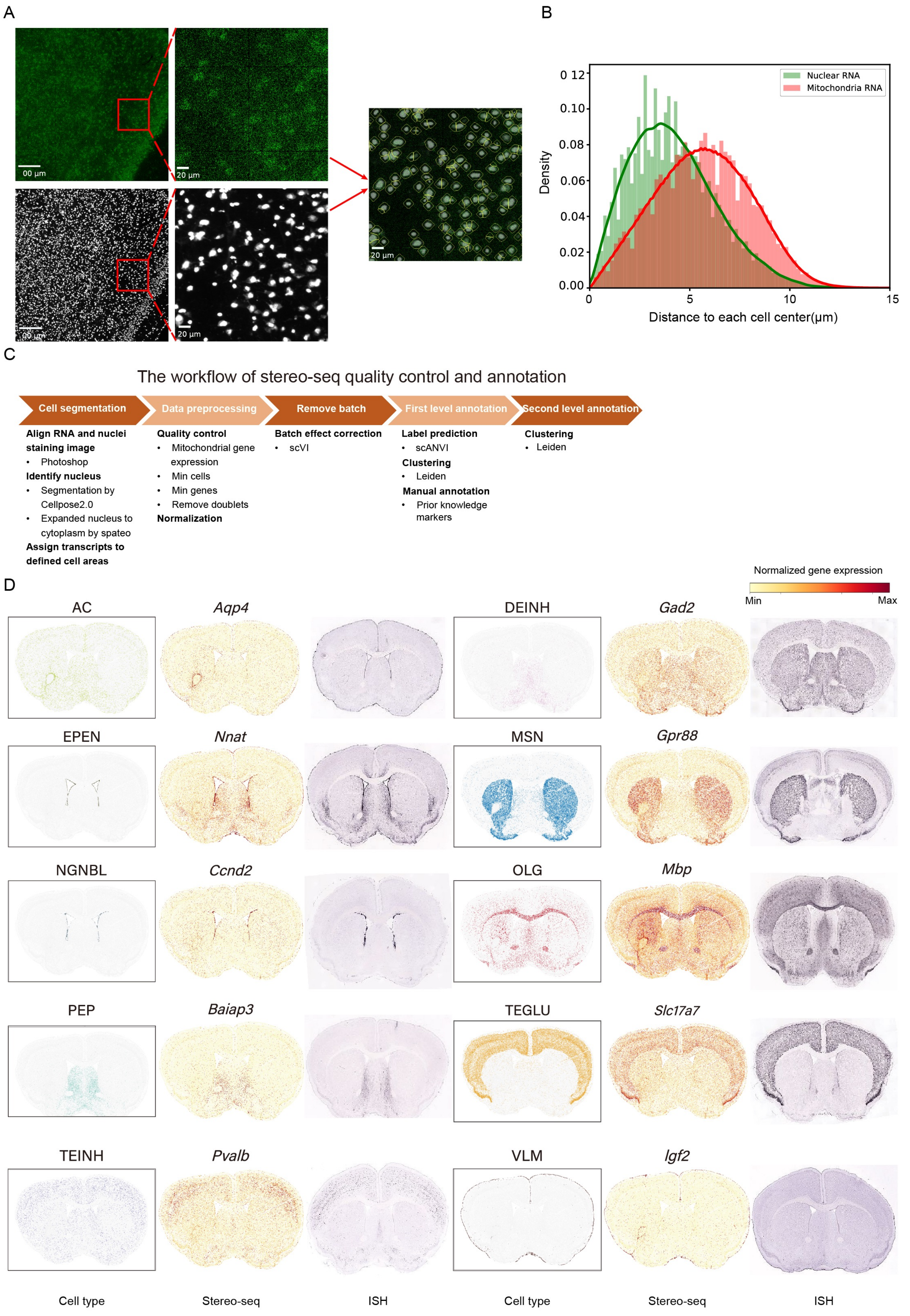
A
00 µm
20 µm
00 µm
20 µm
20 µm
B
Nuclear RNA
Mitochondria RNA
Density
Distance to each cell center(µm)
0 12
0.10
0.08
0.06
0.04
0.02
0.00
0
5
10
15
C
The workflow of stereo-seq quality control and annotation
Cell segmentation
Data preprocessing
Remove batch
First level annotation
Second level annotation
Align RNA and nuclei staining image
• Photoshop
Identify nucleus
• Segmentation by Cellpose2.0
• Expanded nucleus to cytoplasm by spateo
Assign transcripts to defined cell areas
Quality control
• Mitochondrial gene expression
• Min cells
• Min genes
• Remove doublets
Normalization
Batch effect correction
• scVI
Label prediction
• scANVI
Clustering
• Leiden
Manual annotation
• Prior knowledge markers
Clustering
• Leiden
D
Normalized gene expression
Min
Max
AC
Aqp4
DEINH
Gad2
EPEN
Nnat
MSN
Gpr88
NGNBL
Ccnd2
OLG
Mbp
PEP
Baiap3
TEGLU
Slc17a7
TEINH
Pvalb
VLM
Igf2
Cell type
Stereo-seq
ISH
Cell type
Stereo-seq
ISH

**Figure S1 Cell segmentation and quality control workflow, related to Figure 1**

(A) Cell segmentation process: In the upper part, each pixel represents a DNA nanoball (DNB) on the Stereo-seq chip, with 500 nanometers between each DNB. The color intensity indicates the number of RNA transcripts captured per DNB, with darker colors indicating higher transcript counts. The lower part shows cell nuclei stained from the same location, with each white point representing a nucleus. The rightmost image shows the merged DNBs and cell nuclei, illustrating the cell segmentation results. The yellow lines indicate the boundaries of each cell.

(B) The distribution of nuclear-localized RNA (*Malat1* and *Neat1*) and cytoplasmic RNA (mitochondria) in the sampled cells.

(C) Quality control and annotation workflow of Stereo-seq.

(D) Spatial visualization of primary cell types (left) along with corresponding markers (middle) and comparison with Allen Mouse Brain In Situ Hybridization (ISH) data (right).

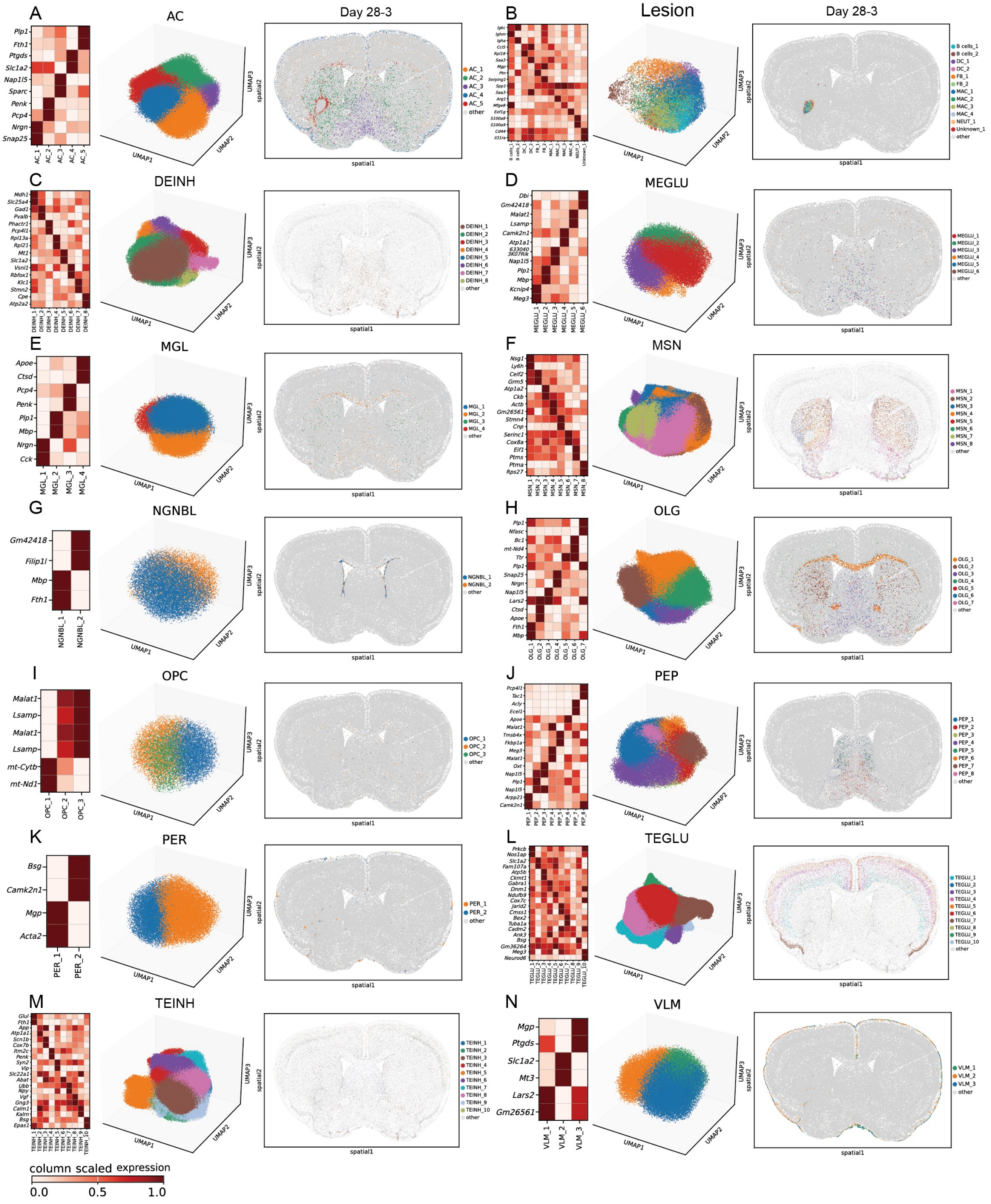

A
AC
Day 28-3
Plp1
Fth1
Ptgds
Slc1a2
Nap1l5
Sparc
Penk
Pcp4
Nrgn
Snap25
AC_1
AC_2
AC_3
AC_4
AC_5
other
B
Lesion
Day 28-3
B cells_1
B cells_2
DC_1
DC_2
FB_1
FB_2
MAC_1
MAC_2
MAC_3
MAC_4
NEUT_1
Unknown_1
other
C
DEINH
DEINH_1
DEINH_2
DEINH_3
DEINH_4
DEINH_5
DEINH_6
DEINH_7
DEINH_8
other
D
MEGLU
Dbi
Gm42418
Malat1
Lsamp
Camk2n1
Atp1a1
Nap1l5
Plp1
Mbp
Kcnip4
Meg3
MEGLU_1
MEGLU_2
MEGLU_3
MEGLU_4
MEGLU_5
MEGLU_6
other
E
MGL
Apoe
Ctsd
Pcp4
Penk
Plp1
Mbp
Nrgn
Cck
MGL_1
MGL_2
MGL_3
MGL_4
other
F
MSN
MSN_1
MSN_2
MSN_3
MSN_4
MSN_5
MSN_6
MSN_7
MSN_8
other
G
NGNBL
Gm42418
Filip1l
Mbp
Fth1
NGNBL_1
NGNBL_2
other
H
OLG
OLG_1
OLG_2
OLG_3
OLG_4
OLG_5
OLG_6
OLG_7
other
I
OPC
Malat1
Lsamp
Malat1
Lsamp
mt-Cytb
mt-Nd1
OPC_1
OPC_2
OPC_3
other
J
PEP
PEP_1
PEP_2
PEP_3
PEP_4
PEP_5
PEP_6
PEP_7
PEP_8
other
K
PER
Bsg
Camk2n1
Mgp
Acta2
PER_1
PER_2
other
L
TEGLU
TEGLU_1
TEGLU_2
TEGLU_3
TEGLU_4
TEGLU_5
TEGLU_6
TEGLU_7
TEGLU_8
TEGLU_9
TEGLU_10
other
M
TEINH
TEINH_1
TEINH_2
TEINH_3
TEINH_4
TEINH_5
TEINH_6
TEINH_7
TEINH_8
TEINH_9
TEINH_10
other
N
VLM
Mgp
Ptgds
Slc1a2
Mt3
Lars2
Gm26561
VLM_1
VLM_2
VLM_3
other
UMAP1
UMAP2
UMAP3
spatial1
spatial2
column scaled expression
0.0
0.5
1.0

**Figure S2 Marker gene expression and UMAP visualization of cell subclasses, related to Figure 1**

(A-N) Average marker gene expression after z-score transformation of cell subclasses (left), UMAP visualization of cell subclasses (middle), spatial visualization of cell subclasses (right).

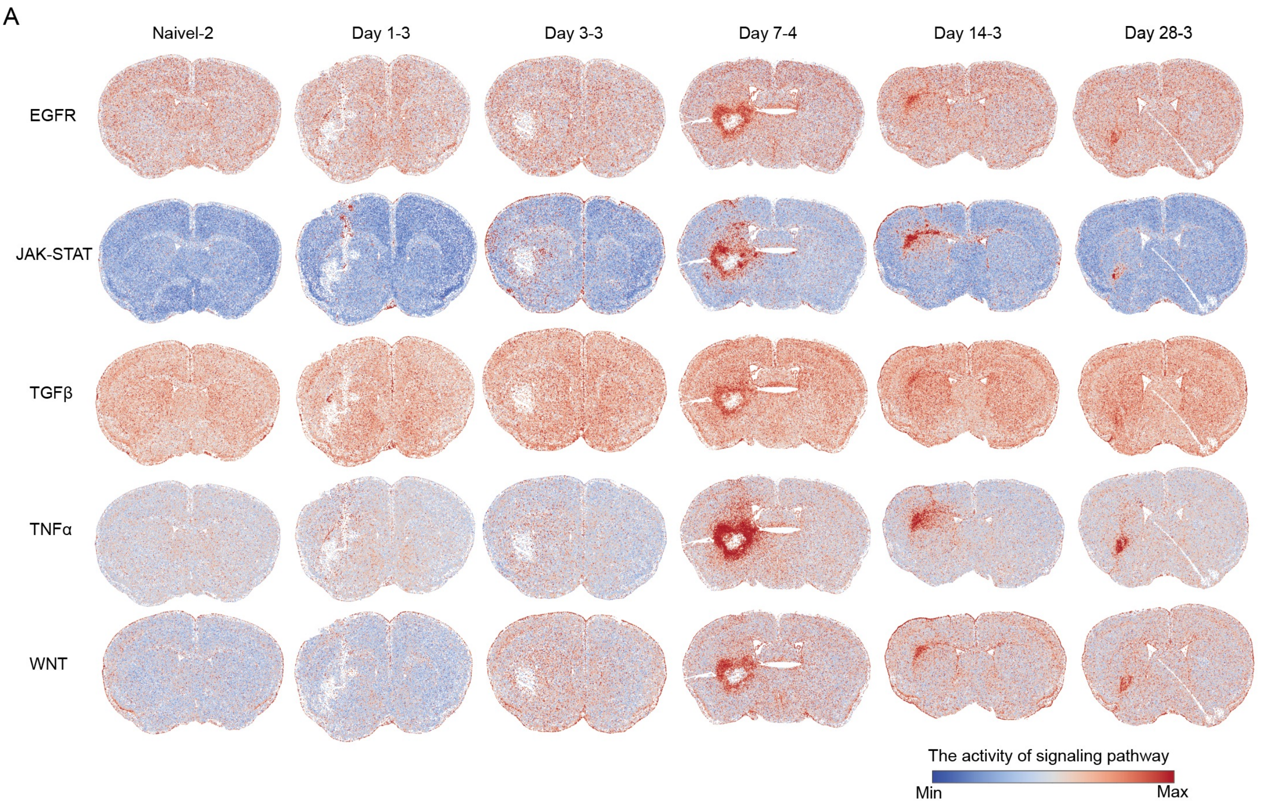

A
Naivel-2
Day 1-3
Day 3-3
Day 7-4
Day 14-3
Day 28-3
EGFR
JAK-STAT
TGFβ
TNFα
WNT
The activity of signaling pathway
Min
Max

**Figure S3 The activity of canonical signaling pathways**

(A) The activity of canonical signaling pathways at various time intervals.

A

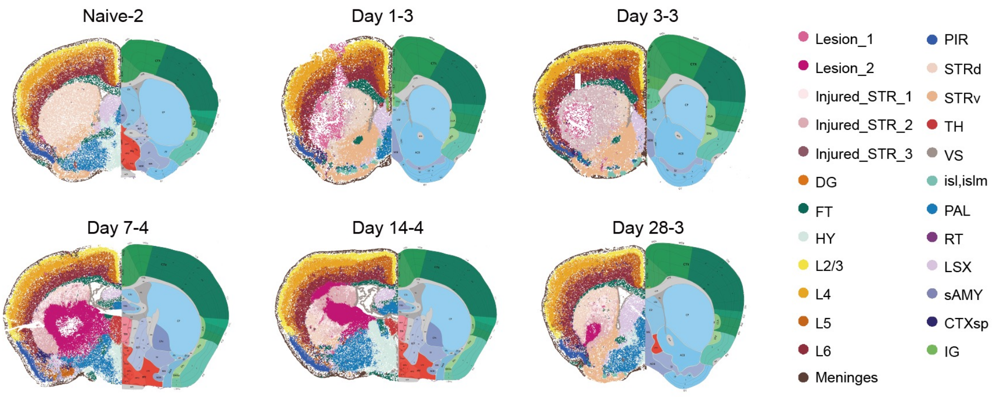

Naive-2
Day 1-3
Day 3-3
Day 7-4
Day 14-4
Day 28-3
Lesion_1
Lesion_2
Injured_STR_1
Injured_STR_2
Injured_STR_3
DG
FT
HY
L2/3
L4
L5
L6
Meninges
PIR
STRd
STRv
TH
VS
isl,islm
PAL
RT
LSX
sAMY
CTXsp
IG


B

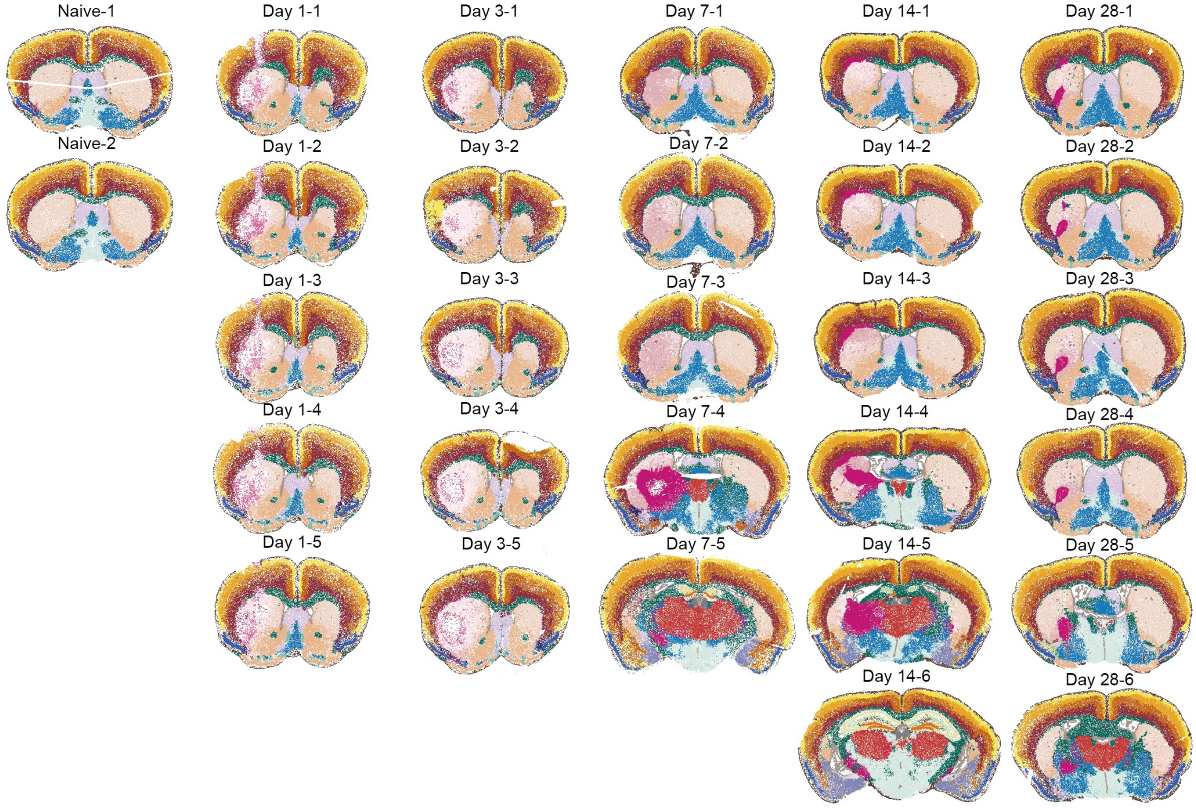

Naive-1
Naive-2
Day 1-1
Day 1-2
Day 1-3
Day 1-4
Day 1-5
Day 3-1
Day 3-2
Day 3-3
Day 3-4
Day 3-5
Day 7-1
Day 7-2
Day 7-3
Day 7-4
Day 7-5
Day 14-1
Day 14-2
Day 14-3
Day 14-4
Day 14-5
Day 14-6
Day 28-1
Day 28-2
Day 28-3
Day 28-4
Day 28-5
Day 28-6

**Figure S4 Anatomical partitioning, related to Figure 2**

(A) The alignment between molecularly defined tissue regions (left, colored based on molecular tissue region identities) and anatomically defined tissue regions (right, colored based on anatomically defined tissue regions). Each dot in the left hemisphere represents a cell. The anatomical definition of the right hemisphere comes from the Allen Brain Atlas. The abbreviation of the tissue region is consistent with the Allen Mouse Brain Reference Map. L2/3, cortical layer 2 and cortical layer 3; L4, cortical layer 4; L5, cortical layer 5; L6, cortical layer 6; lesion_1, lesion 1; lesion_2, lesion 2; Injured_STR_1, injured striatum 1; Injured_STR_2, injured striatum 2; Injured_STR_3, injured striatum 3; DG, dentate gyrus; FT, fiber tracts; HY, hypothalamus; PIR, piriform area; STRd, striatum dorsal region; STRv, striatum ventral region; TH, thalamus; VS, ventricular systems; isl,islm, islands of calleja and major island of calleja; PAL, pallidum. RT, reticular nucleus of the thalamus; LSX, lateral septal complex; sAMY, striatum-like amygdalar nuclei; CTXsp, cortical subplate;IG,Induseum griseum.

(B) Molecularly defined tissue regions across all slices, colored by defined regions.

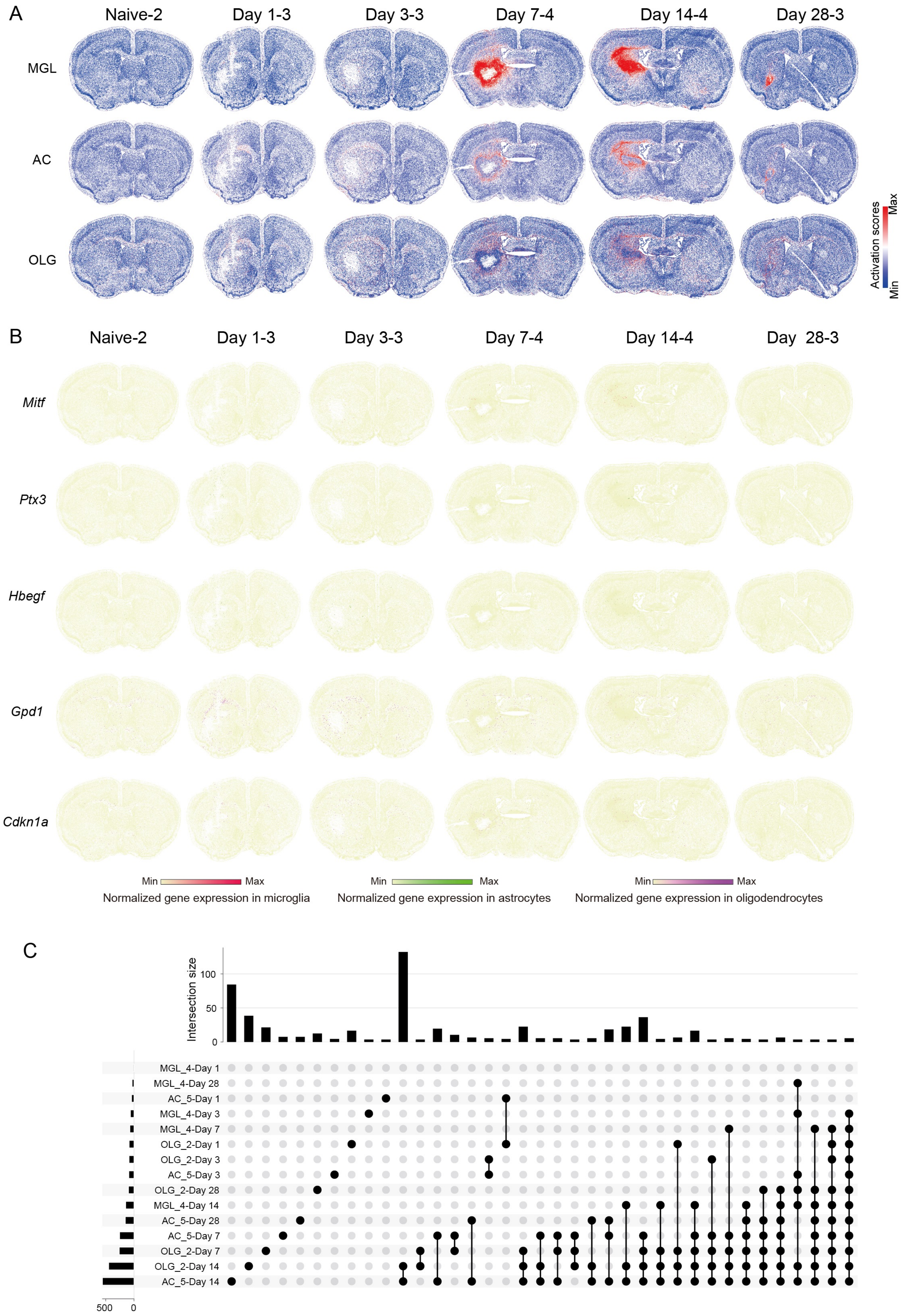

A
Naive-2
Day 1-3
Day 3-3
Day 7-4
Day 14-4
Day 28-3
MGL
AC
OLG
Activation scores
Max
Min
B
Naive-2
Day 1-3
Day 3-3
Day 7-4
Day 14-4
Day 28-3
Mitf
Ptx3
Hbegf
Gpd1
Cdkn1a
Min Max
Normalized gene expression in microglia
Min Max
Normalized gene expression in astrocytes
Min Max
Normalized gene expression in oligodendrocytes
C
Intersection size
100
50
0
MGL_4-Day 1
MGL_4-Day 28
AC_5-Day 1
MGL_4-Day 3
MGL_4-Day 7
OLG_2-Day 1
OLG_2-Day 3
AC_5-Day 3
OLG_2-Day 28
MGL_4-Day 14
AC_5-Day 28
AC_5-Day 7
OLG_2-Day 7
OLG_2-Day 14
AC_5-Day 14
500
0

**Figure S5 Spatial distribution and transcriptional signatures of glial activation, related to Figure 3**

(A) Spatial visualization of activation scores across different timepoints post-injury, for three types of glial cells: microglia (MGL), astrocytes (AC), and oligodendrocytes (OLG). The color intensity corresponds to the degree of activation, with red indicating the highest activation score and blue representing the lowest.

(B) Spatial visualization of genes specifically activated in three types of glial cells. Each plot illustrates the spatial distribution of select genes within their respective glial cell types at designated time points. Background cells are represented in yellow to highlight the gene expression in specific glial cell type. Specifically, *Mitf* is visualized in microglia, *Ptx3* and *Hbegf* in astrocytes, Gpd1 and *Cdkn1a* in oligodendrocytes.

(C) UpSet plot delineates the overlap of differentially expressed genes across three types of activated glial cells at various timepoints. The differentially expressed genes were identified by comparing each activated glial cell subclasses at each timepoint to its corresponding naive state. The horizontal bar chart on the bottom left displays the total number of differentially expressed genes within each dataset, while the vertical bar chart on the upper right illustrates the intersections between these datasets (with black dots indicating the presence of an intersection).

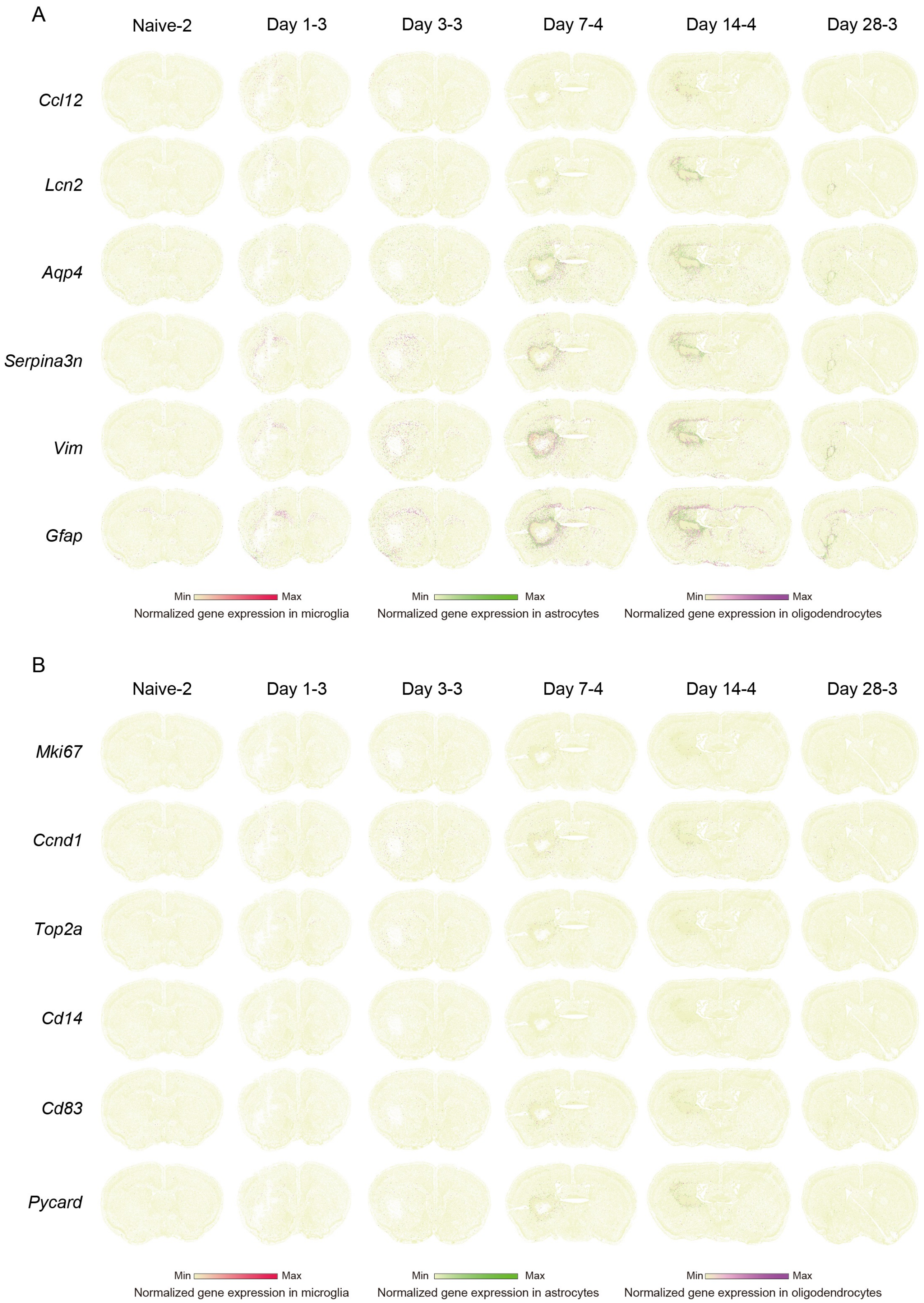
A
Naive-2
Day 1-3
Day 3-3
Day 7-4
Day 14-4
Day 28-3
Ccl12
Lcn2
Aqp4
Serpina3n
Vim
Gfap
Min
Max
Normalized gene expression in microglia
Min
Max
Normalized gene expression in astrocytes
Min
Max
Normalized gene expression in oligodendrocytes
B
Naive-2
Day 1-3
Day 3-3
Day 7-4
Day 14-4
Day 28-3
Mki67
Ccnd1
Top2a
Cd14
Cd83
Pycard
Min
Max
Normalized gene expression in microglia
Min
Max
Normalized gene expression in astrocytes
Min
Max
Normalized gene expression in oligodendrocytes

**Figure S6 Transcriptional signatures of glial activation, related to Figure 3**

(A) Spatial visualization of co-activated genes across three activated glial cells. Background cells are represented in yellow to highlight the gene expression in three glial cell type.

(B) Spatial visualization of genes that are upregulated specifically in astrocytes and oligodendrocytes. Changes in gene expression levels are shown for only these two cell types, with the other cell types showing the background in yellow.

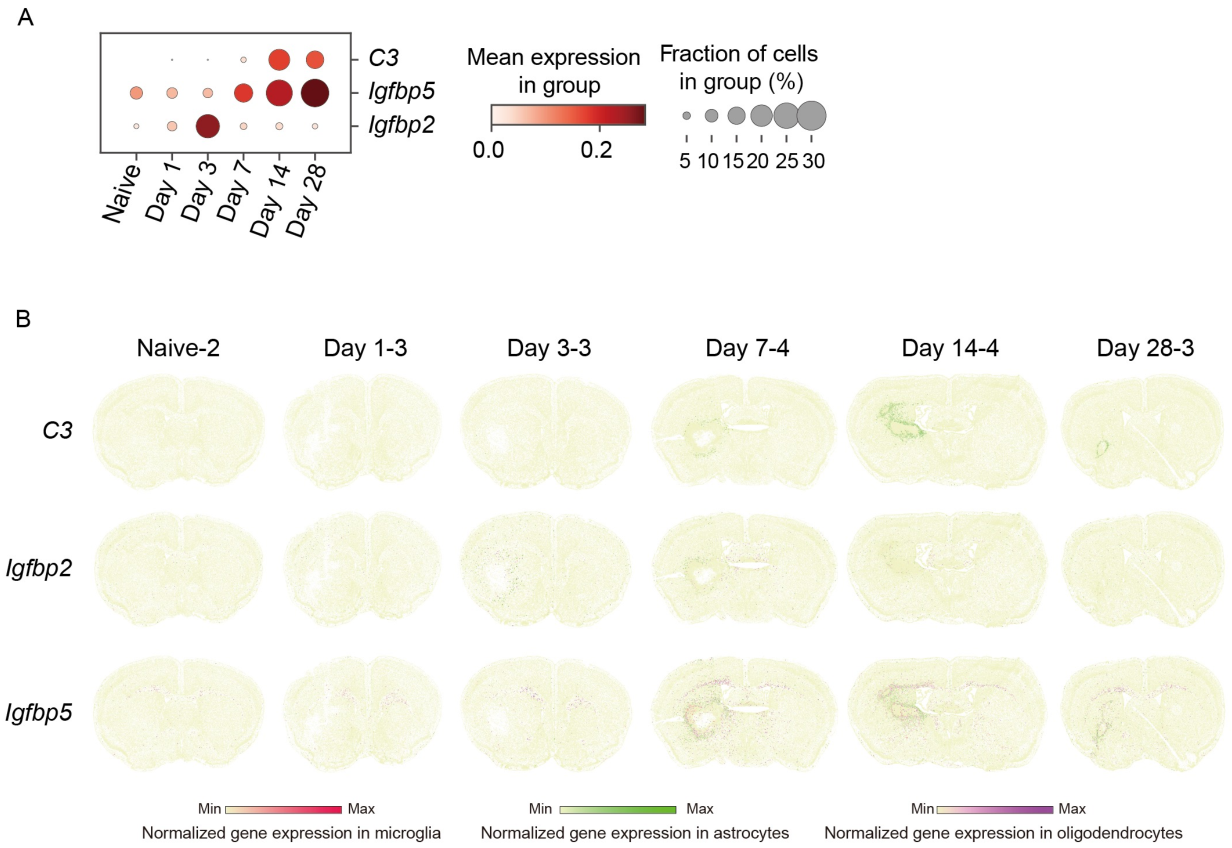

A
C3
Igfbp5
Igfbp2
Naive
Day 1
Day 3
Day 7
Day 14
Day 28
Mean expression in group
0.0
0.2
Fraction of cells in group (%)
5 10 15 20 25 30
B
Naive-2
Day 1-3
Day 3-3
Day 7-4
Day 14-4
Day 28-3
C3
Igfbp2
Igfbp5
Min
Max
Normalized gene expression in microglia
Min
Max
Normalized gene expression in astrocytes
Min
Max
Normalized gene expression in oligodendrocytes

**Figure S7 Functional diversity of astrocytes, related to Figure 3**

(A) Dot plot depicts the temporal expression patterns of genes *C3*, *Igfbp5*, and *Igfbp2* in activated astrocytes (AC_5) from the naive state through various post-ICH stages (Day 1 to Day 28). The color intensity represents the mean expression level within the group, while the dot size indicates the percentage of cells expressing the gene in the group.

(B) Spatial visualization of *C3*, *Igfbp5*, and *Igfbp2,* color by gene expression level.

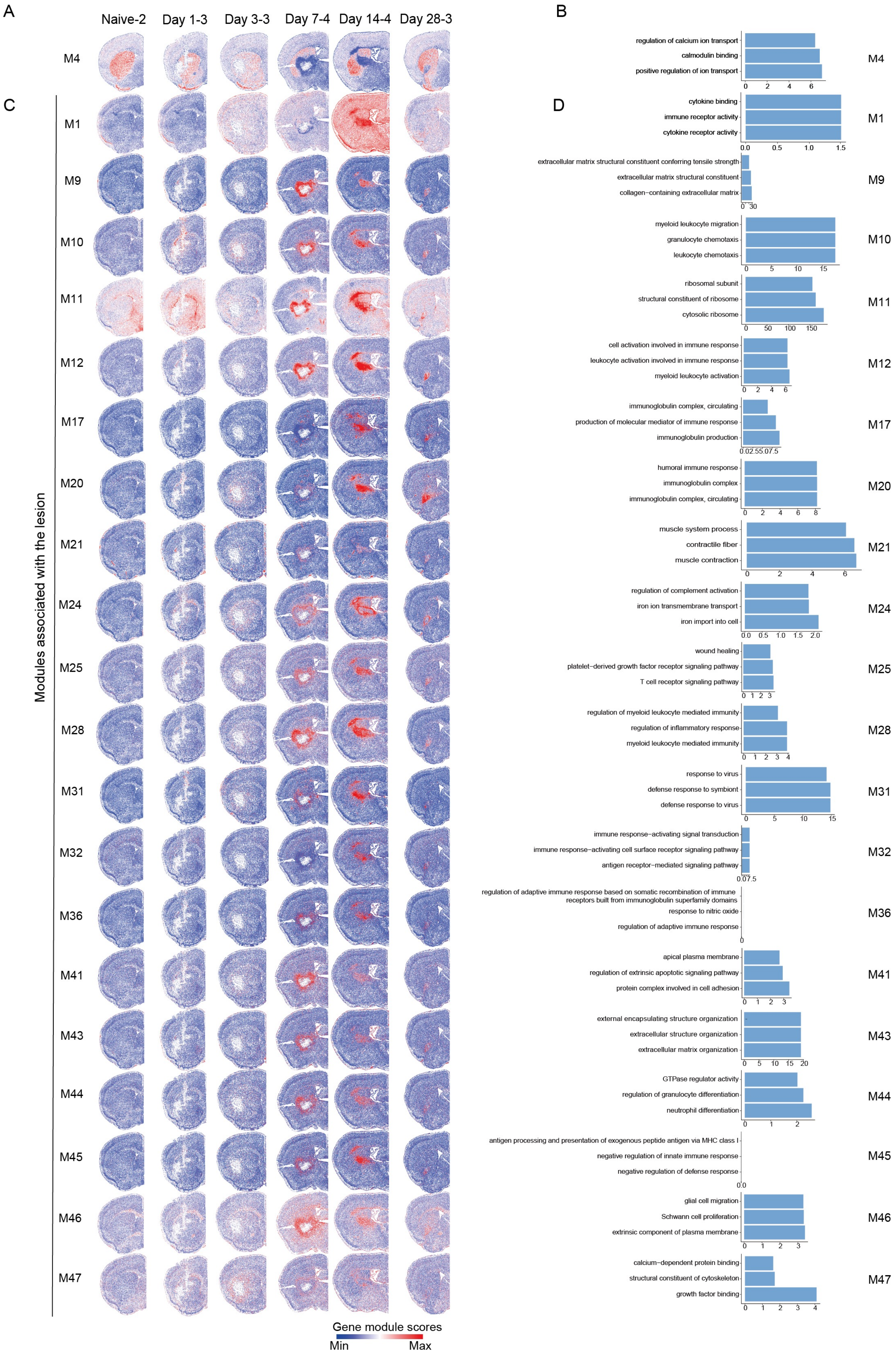
A
Naive-2
Day 1-3
Day 3-3
Day 7-4
Day 14-4
Day 28-3
M4
C
Modules associated with the lesion
M1
M9
M10
M11
M12
M17
M20
M21
M24
M25
M28
M31
M32
M36
M41
M43
M44
M45
M46
M47
Gene module scores
Min
Max
B
regulation of calcium ion transport
calmodulin binding
positive regulation of ion transport
D
cytokine binding
immune receptor activity
cytokine receptor activity
extracellular matrix structural constituent conferring tensile strength
extracellular matrix structural constituent
collagen−containing extracellular matrix
myeloid leukocyte migration
granulocyte chemotaxis
leukocyte chemotaxis
ribosomal subunit
structural constituent of ribosome
cytosolic ribosome
cell activation involved in immune response
leukocyte activation involved in immune response
myeloid leukocyte activation
immunoglobulin complex, circulating
production of molecular mediator of immune response
immunoglobulin production
humoral immune response
immunoglobulin complex
immunoglobulin complex, circulating
muscle system process
contractile fiber
muscle contraction
regulation of complement activation
iron ion transmembrane transport
iron import into cell
wound healing
platelet−derived growth factor receptor signaling pathway
T cell receptor signaling pathway
regulation of myeloid leukocyte mediated immunity
regulation of inflammatory response
myeloid leukocyte mediated immunity
response to virus
defense response to symbiont
defense response to virus
immune response−activating signal transduction
immune response−activating cell surface receptor signaling pathway
antigen receptor−mediated signaling pathway
regulation of adaptive immune response based on somatic recombination of immune receptors built from immunoglobulin superfamily domains
response to nitric oxide
regulation of adaptive immune response
apical plasma membrane
regulation of extrinsic apoptotic signaling pathway
protein complex involved in cell adhesion
external encapsulating structure organization
extracellular structure organization
extracellular matrix organization
GTPase regulator activity
regulation of granulocyte differentiation
neutrophil differentiation
antigen processing and presentation of exogenous peptide antigen via MHC class I
negative regulation of innate immune response
negative regulation of defense response
glial cell migration
Schwann cell proliferation
extrinsic component of plasma membrane
calcium−dependent protein binding
structural constituent of cytoskeleton
growth factor binding

**Figure S8 Additional gene modules, related to Figure 4**

(A) & (C) Spatial visualization of gene modules, color by module scores. (A) delineates gene module within normal function, and (C) focuses on gene modules with lesion. Key: 'M' denotes modules.

(B) & (D) Gene ontology (go) enrichment analysis of gene modules from (A) & (C). X-axis: –log10 (p-adjust) score; Y-axis: GO terms. Only top 3 terms (p-adjust < 0.01) are listed for each gene module.

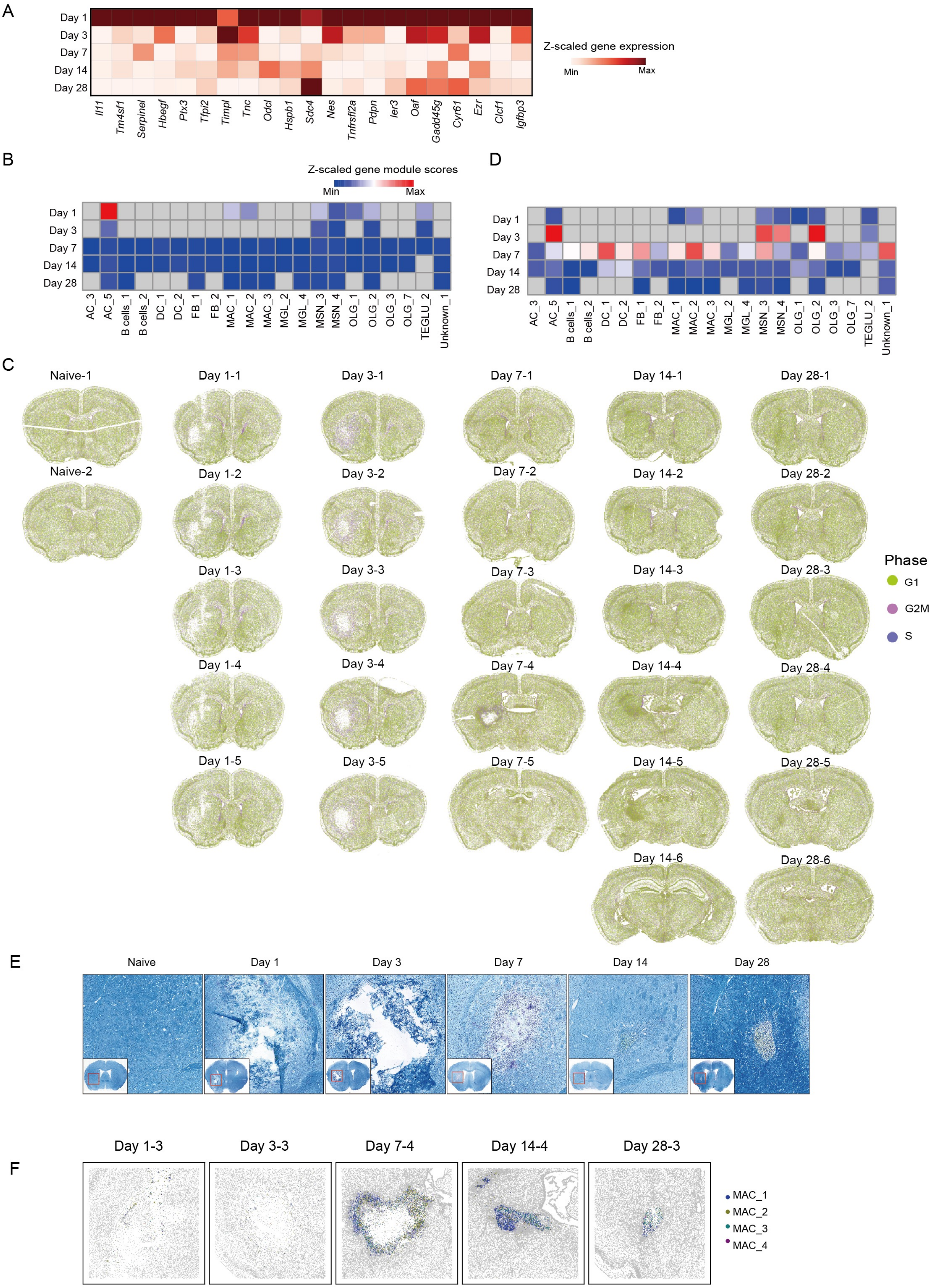

A
Day 1
Day 3
Day 7
Day 14
Day 28
Z-scaled gene expression
Min
Max
Il11
Tm4sf1
Serpinel
Hbegf
Ptx3
Tfpi2
Timpl
Tnc
Odcl
Hspb1
Sdc4
Nes
Tnfrsfl2a
Pdpn
Ier3
Oaf
Gadd45g
Cyr61
Ezr
Clcf1
Igfbp3
B
Z-scaled gene module scores
Min
Max
AC_3
AC_5
B cells_1
B cells_2
DC_1
DC_2
FB_1
FB_2
MAC_1
MAC_2
MAC_3
MGL_2
MGL_4
MSN_3
MSN_4
OLG_1
OLG_2
OLG_3
OLG_7
TEGLU_2
Unknown_1
D
C
Naive-1
Naive-2
Day 1-1
Day 1-2
Day 1-3
Day 1-4
Day 1-5
Day 3-1
Day 3-2
Day 3-3
Day 3-4
Day 3-5
Day 7-1
Day 7-2
Day 7-3
Day 7-4
Day 7-5
Day 14-1
Day 14-2
Day 14-3
Day 14-4
Day 14-5
Day 14-6
Day 28-1
Day 28-2
Day 28-3
Day 28-4
Day 28-5
Day 28-6
Phase
G1
G2M
S
E
Naive
Day 1
Day 3
Day 7
Day 14
Day 28
F
Day 1-3
Day 3-3
Day 7-4
Day 14-4
Day 28-3
MAC_1
MAC_2
MAC_3
MAC_4

**Figure S9 Temporal characteristics of the lesion, related to Figure 4**

(A) Heatmap depicting the average expression of genes of gene module 42 across different time points within the lesion.

(B) Heatmap illustrating the scores of different cell subclasses from gene module 42 at different time points within the lesion.

(C)The distribution of cell cycle phases within all cells at various timepoints, from the naive state through subsequent days post-ICH (day 1 to day 28).

(D) Similar to (B), but for gene module 37.

(E) CV/LFB stained sections at different times.

(F) Spatial visualization of macrophage subclasses in the lesion.

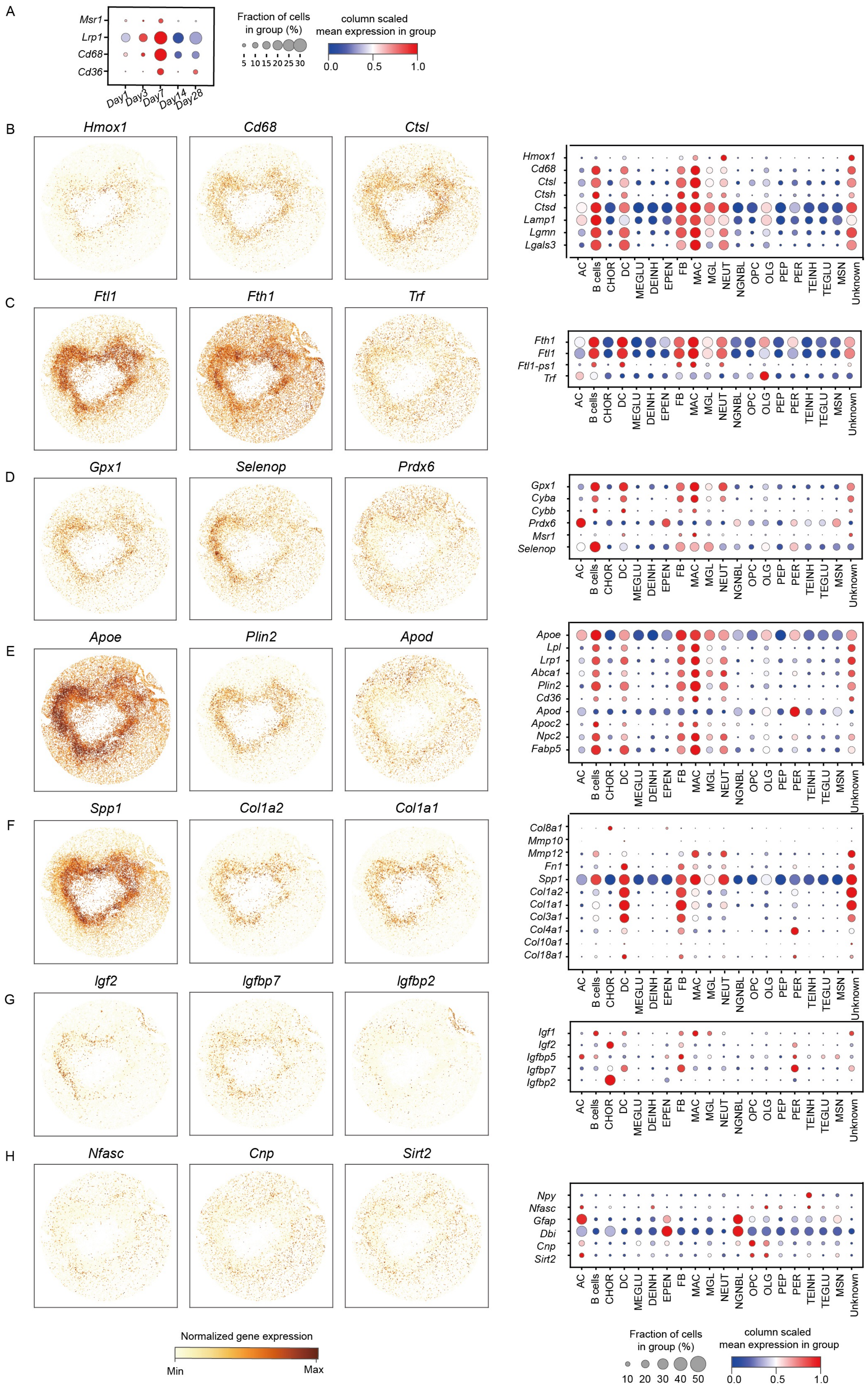
A
Msr1
Lrp1
Cd68
Cd36
Day1
Day3
Day7
Day14
Day28
Fraction of cells in group (%)
5 10 15 20 25 30
column scaled mean expression in group
0.0 0.5 1.0
B
Hmox1
Cd68
Ctsl
Hmox1
Cd68
Ctsl
Ctsh
Ctsd
Lamp1
Lgmn
Lgals3
C
Ftl1
Fth1
Trf
Fth1
Ftl1
Ftl1-ps1
Trf
D
Gpx1
Selenop
Prdx6
Gpx1
Cyba
Cybb
Prdx6
Msr1
Selenop
E
Apoe
Plin2
Apod
Apoe
Lpl
Lrp1
Abca1
Plin2
Cd36
Apod
Apoc2
Npc2
Fabp5
F
Spp1
Col1a2
Col1a1
Col8a1
Mmp10
Mmp12
Fn1
Spp1
Col1a2
Col1a1
Col3a1
Col4a1
Col10a1
Col18a1
G
Igf2
Igfbp7
Igfbp2
Igf1
Igf2
Igfbp5
Igfbp7
Igfbp2
H
Nfasc
Cnp
Sirt2
Npy
Nfasc
Gfap
Dbi
Cnp
Sirt2
AC
B cells
CHOR
DC
MEGLU
DEINH
EPEN
FB
MAC
MGL
NEUT
NGNBL
OPC
OLG
PEP
PER
TEINH
TEGLU
MSN
Unknown
Normalized gene expression
Min
Max
Fraction of cells in group (%)
10 20 30 40 50
column scaled mean expression in group
0.0 0.5 1.0

**Figure S10 Visualization of genes related to the pathological process of intracerebral hemorrhage, related to Figure 5**

(A) The mean expression of scavenger receptor in different time points within the lesion.

(B) The left side of the figure presents spatial gene expression maps of phagocytosis-related genes (*Hmox1*, *Cd68*, *Ctsl*) within the lesion on day 7 post-ICH. The right side of the figure displays the mean expression of these phagocytosis-related genes in different cell types within the lesion on day 7 post-ICH. Additionally, genes with various functions are shown in (C) Iron metabolism, (D) Oxidation and antioxidation, (E) Lipid metabolism and cholesterol transport, (F) Extracellular matrix, (G) Growth factor, (H) Neural repair.

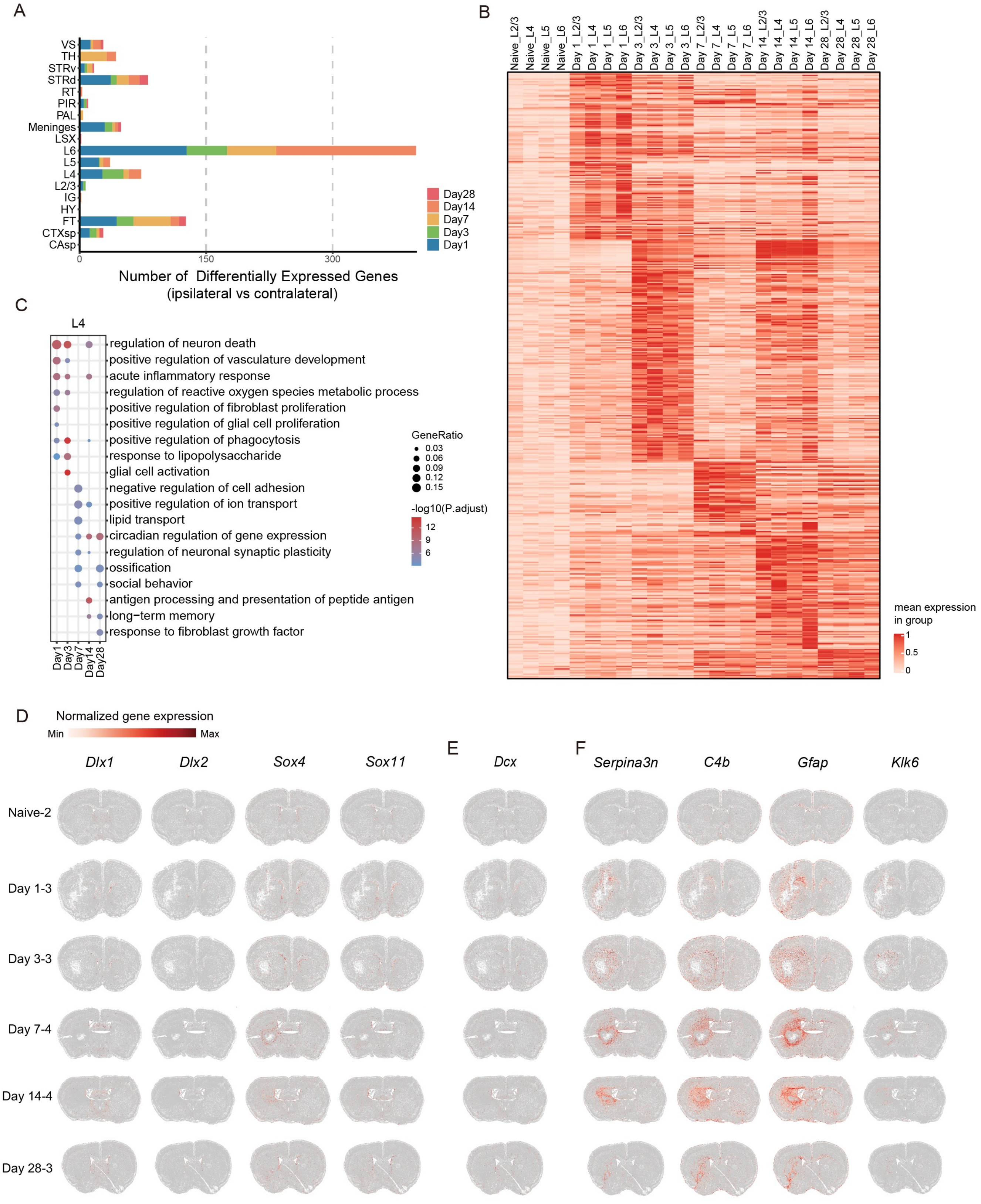

A
VS
TH
STRv
STRd
RT
PIR
PAL
Meninges
LSX
L6
L5
L4
L2/3
IG
HY
FT
CTXsp
CAsp
0
150
300
Number of Differentially Expressed Genes
(ipsilateral vs contralateral)
Day28
Day14
Day7
Day3
Day1
B
Naive_L2/3
Naive_L4
Naive_L5
Naive_L6
Day 1_L2/3
Day 1_L4
Day 1_L5
Day 1_L6
Day 3_L2/3
Day 3_L4
Day 3_L5
Day 3_L6
Day 7_L2/3
Day 7_L4
Day 7_L5
Day 7_L6
Day 14_L2/3
Day 14_L4
Day 14_L5
Day 14_L6
Day 28_L2/3
Day 28_L4
Day 28_L5
Day 28_L6
mean expression
in group
1
0.5
0
C
L4
regulation of neuron death
positive regulation of vasculature development
acute inflammatory response
regulation of reactive oxygen species metabolic process
positive regulation of fibroblast proliferation
positive regulation of glial cell proliferation
positive regulation of phagocytosis
response to lipopolysaccharide
glial cell activation
negative regulation of cell adhesion
positive regulation of ion transport
lipid transport
circadian regulation of gene expression
regulation of neuronal synaptic plasticity
ossification
social behavior
antigen processing and presentation of peptide antigen
long−term memory
response to fibroblast growth factor
Day1
Day3
Day7
Day14
Day28
GeneRatio
0.03
0.06
0.09
0.12
0.15
-log10(P.adjust)
12
9
6
D
Normalized gene expression
Min
Max
Dlx1
Dlx2
Sox4
Sox11
Naive-2
Day 1-3
Day 3-3
Day 7-4
Day 14-4
Day 28-3
E
Dcx
F
Serpina3n
C4b
Gfap
Klk6

**Figure S11 Molecular changes in different brain regions after intracerebral hemorrhage, related to Figure 6**

(A) The number of differentially expressed genes in various ipsilateral brain regions at different time points following ICH compared to the contralateral group. Significant genes were identified based on specific criteria: logfoldchange≥1, scores ＞ 5, and p-adjust＜0.001.

(B) The heatmap illustrates the average expression levels of genes that are differentially expressed across cortical regions at various time points after ICH, in comparison to naïve controls. Columns represent distinct cortical areas at specific time points post-ICH, while rows correspond to individual genes.

(C) Bubble plots depicts the GO enrichment analysis results for genes differentially expressed in the L4 cortical layer at successive time points following ICH.

(D) Spatial visualization of *Dlx1, Dlx2, Sox4, and Sox11*.

(E) Spatial visualization of *Dcx*.

(F) Spatial visualization of *Serpina3n, C4b, Gfap, and Klk6*.
